# Multiscale Responsive Kinetic Modeling: Quantifying Biomolecular Reaction Flux under Varying Electrochemical Conditions

**DOI:** 10.1101/2024.08.01.606205

**Authors:** Hannah Weckel-Dahman, Ryan Carlsen, Jessica M.J. Swanson

## Abstract

Attaining a complete thermodynamic and kinetic characterization for processes involving multiple interconnected rare-event transitions remains a central challenge in molecular biophysics. This challenge is amplified when the process must be understood under a range of reaction conditions. Herein, we present a condition-responsive kinetic modeling framework that can combine the strengths of bottom-up rate quantification from multiscale simulations with top-down solution refinement using experimental data. Although this framework can be applied to any process, we demonstrate its use for electrochemically driven transport through channels and transporters. Using the Cl^−^ /H^+^ antiporter ClC-ec1 as a model system, we show how robust and predictive kinetic solutions can be obtained when the solution space is grounded by thermodynamic constraints, seeded through multiscale rate quantification, and further refined with experimental data, such as electrophysiology assays. Turning to the Shaker K^+^ channel, we demonstrate that robust solutions and biophysical insights can also be obtained with sufficient experimental data. This multi-pathway method proves capable of identifying single-pathway dominant mechanisms but also highlights that competing and off-pathway flux is still essential to replicate experimental findings and to describe concentration-dependent channel rectification.

## 1. INTRODUCTION

The mechanisms of most biomolecular processes are well described by reactant and product states that are separated by a finite number of metastable intermediates (states) connected by rare-event transitions. As long as rates of transitions are known, the reactive flux through these intermediates in response to non-equilibrium starting populations is easily described by a set of ordinary differential equations according to the kinetic master equation. Connecting this coarse-grained kinetic description with a molecular-level understanding of the involved rare-event transition ensembles lends unprecedented insight into the mechanism and how it responds to non-equilibrium conditions. However, defining a robust and accurate kinetic model is challenging as it requires identifying all relevant intermediates, determining which intermediates are connected by kinetically relevant transitions, and quantifying the transition rates for the range of conditions of interest. To help address these challenges, we present multiscale responsive kinetic modeling (MsRKM), a framework that combines the strengths of computational characterization of intermediates and rare-event transitions with experimental data to identify robust kinetic solutions. This enables the description of any biomolecular processes involving multiple rare event transitions in terms of the underlying mechanistic reaction network and the ensemble flux through intermediates in response to initial nonequilibrium and equilibrium conditions. MsRKM includes the dynamic response of the underlying rate coefficients to the influence of transmembrane voltage, enabling the description of electrochemically driven transport through channels and transporters.

Part of the motivation for this work is an increasing recognition of the importance of kinetic selection and mechanistic heterogeneity.^3-9^ Although some processes are certainly dominated by a single pathway (sequence of intermediates), many others involve off-pathway transitions and condition-varying flux through alternative pathways that can play essential roles in mechanistic outcomes (Figure 1). A prime example is kinetic proofreading, as originally envisioned by Hopefield^10^ and Ninio^11^, in which competing pathways increase substrate selectivity when a slow step follows substrate association. This decreases the error rates in processes requiring high fidelity, such as replication, translation, and transcription^12^, and has also been demonstrated in signal transduction^13^ and pathogen recognition^14, 15^. Kinetic selection can also enable condition-specific mechanistic outcomes, as demonstrated in molecular machines that transform input-free energy (e.g., from an abundance of ATP) into various cellular processes. For example, branched networks play a central role in explaining both chemomechanical coupling^16^ and the different mechanistic steps observed in motor proteins such as myosin and kinesin under different ATP/ADP ratios ^17, 18^.

**Figure 1.**
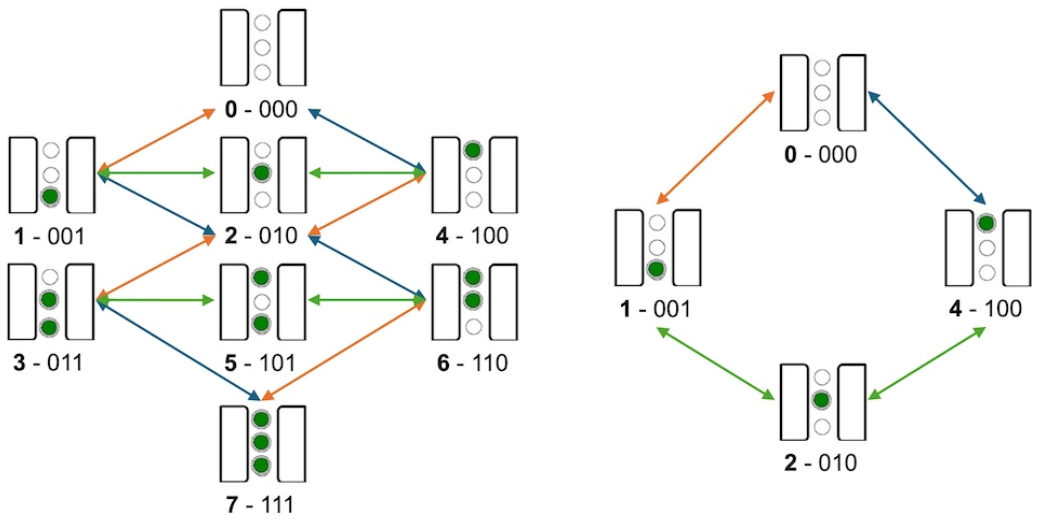
Full reaction network for channel with 3 ion binding sites (left) compared to single-pathway simplification of ion transport process (right). Intermediates are labeled with both a binary description of the bound ions with matching decimal label and state image.

Another domain where kinetic selection is proving important is the controlled transport of solutes across membranes. Membrane channels and transporters play pivotal roles in a myriad of cellular processes (maintaining concentration gradients, triggering action potentials or cell signaling, importing nutrients, exporting waste and toxins, and regulating intracellular signaling) through the translocation of ions and substrates across cellular membranes. In transporters for which consistent stoichiometry must be obeyed across a range of conditions, such as the Na^+^/K^+^ antiporter, exact precision via a single transport pathway may indeed be important. For others, however, kinetic selection between competing pathways explains non-integral and variable stoichiometries and enables essential function. For example, the sodium/glucose transporter was shown to use a substrate slip pathway to exchange some degree of efficiency for toxin discrimination.^8, 19^ Additionally, the multidrug efflux pump EmrE was shown to employ multiple transport pathways to enable different transport regimes, including symport, antiport, and uniport, under different electrochemical driving forces.^20^ These two examples demonstrate how the electrochemical conditions can be a controlling factor in substrate transport, as opposed to the more commonly recognized ligand binding, voltage gating, or ATP hydrolysis. Collectively, these findings demonstrate the importance of modeling transporters with a network description that includes the influences of competing pathways. However, it remains an outstanding question if ion channels also involve competing transport pathways to the degree that a network description is essential to understanding their transport mechanisms. The methods described herein were designed to help address this question.

Kinetic modeling can generally be approached in a ‘top-down’ physics-based theoretical manner using system knowledge and experimental data or in a ‘bottom-up’ tour-de-force quantification of rate coefficients for every transition, typically with simulations. Markov state modeling has become a powerful approach of the latter form, stitching together simulation data to define metastable states and quantifying their transition probabilities.^21, 22^ MsRKM can be used in any combination of purely top-down, as demonstrated for the Shaker Kv channel herein, or purely bottom-up, using simulation-based rate coefficients to describe system behavior under non-equilibrium conditions and test their ability to replicate experimental data. However, it is more powerfully employed by combining the two—narrowing the kinetic solution space based on calculated rate coefficients and refining the solution based on experimental data. This enables an experimentally driven theoretical framework that iteratively tests for consistency between molecular-level characterization and experimental behavior.

The choice of the word ‘responsive’ in the acronym deserves explanation. The ability of a kinetic model to describe a range of conditions requires the underlying rate coefficients to adapt to the conditions of interest. Since unimolecular rate coefficients typically remain stable despite changes in reactant and product concentrations, a single set of rate coefficients can generally capture non-equilibrium conditions by accommodating shifts in concentrations with bimolecular uptake coefficients. In contrast, rate coefficients can shift significantly in response to pH and voltage. MsRKM was developed to explicitly ‘respond to’ changes in voltage with voltage-dependent rate coefficients. This does not, however, include large voltage-induced conformational changes such as those seen in voltage-gated channels. Nor does the current implementation account for significant changes due to pH, which are often associated with a conformational change. Rather, distinct kinetic solutions should be developed for significantly different conformational ensembles. Future efforts will focus on an explicit inclusion of pH-dependent conformational changes and the corresponding rate changes. With the inclusion of voltage-dependent rate coefficients, processes driven by electrochemical gradients can be described by a single set of base (Δ Ψ= 0mV) rate coefficients, capturing non-equilibrium flux under varying electrochemical driving forces. This enables a direct comparison to typical electrophysiology assays.

MsRKM is demonstrated on two membrane-bound proteins: ClC-ec1, a secondary-active exchanger, and the Shaker Kv channel. ClC-ec1 is a well-studied Cl^−^/H^+^ antiporter from Escherichia coli that is involved in cellular ion homeostasis and plays an important, though incompletely understood, role in the cell’s acid resistance mechanism.^23, 24^ Structurally, ClC-ec1 operates as a homodimer, with each subunit containing an independent ion-conducting pore.^25^ Based on its measured reversal potential, ClC-ec1 was shown to exchange ions with a 2.2 Cl^−^ to 1 H^+^ stoichiometry.^26^ This exchange is influenced by pH-dependent conformational changes near and within the pore^26-29^. Structure-function studies have identified the key residues involved in ion transport, and electrophysiology assays have revealed a range current-voltage behaviors under different electrochemical conditions.^24-26, 29-32^ Simulations have characterized the proton transport mechanisms^33, 34^ and pH-dependent conformational changes^27-29^. Additionally, previous kinetic models have suggested that ClC-ec1 uses multiple ion exchange pathways, each contributing to the total flux with varying magnitudes under different pH conditions.^6, 9, 31^ Collectively, the extensive experimental and simulation-based analyses make ClC-ec1 an ideal, though still very challenging, model system for MsRKM—testing the method’s ability to capture nuanced transporter behavior in response to changing electrochemical conditions.

Shaker is a voltage-gated potassium channel originally found in *Drosophila funebris*^35^ that is essential for repolarizing the neuronal membrane after an action potential, thereby regulating neuronal excitability and firing patterns.^36, 37^ Shaker is comprised of four subunits that form a central pore through which potassium ions pass when unobstructed by the activation gate.^38, 39^ While Shaker has two distinct inactivation modes,^40^ we focus on the fully open channel. Several mechanisms have been proposed for Shaker that vary in ion occupancy, the order of uptake and release,^41-44^ and the presence or absence of water in the selectivity filter.^45^ However, mechanistic cycles proposed for the open-conformation of Shaker are primarily represented by single-pathway dominant mechanism regardless of the experimental conditions tested, though there is debate on the number of cycles present for homologous proteins.^43, 46^ Additionally, two interesting studies using molecular dynamics simulations run with applied voltages and combined with ion binding graph representations showed many contributing pathways and a mechanistic shift between those observed under chemical gradients and voltages.^47, 48^ However, the applied voltages were far larger than biologically or experimentally relevant magnitudes, which perhaps explains why the findings have not been followed up on. Collectively, these facets make Shaker ideal for testing ground for MsRKM—particularly testing if a full network kinetic characterization will retain the presumed mechanistic insensitivity to electrochemical conditions and if it supports a mechanism dominated by a single mechanistic pathway or if multiple pathways contribute.

For ClC-ec1 we find that a combined bottom-up/top-down approach integrating sufficient simulation and experimental data in MsRKM produces robust kinetic models that are predictive of experimental behavior across a range of electrochemical conditions. For Shaker, MsRKM identifies robust solutions using a purely top-down approach integrating increasing types of experimental data and biophysical insight. Oversimplification of the involved intermediates proves incapable of reproducing all experimental trends, demonstrating how the method can help refine flawed network descriptions. Interestingly, through continued solution refinement MsRKM iteratively rules out many multi-pathway solutions to converge on a single-pathway dominant mechanism that does not vary significantly under electrochemical conditions. Yet even with this mechanism, off pathway flux and a network description of the Shaker channel proves essential for replicating experimental observations.

## 2. THEORY AND METHODS

The essential ingredients for any kinetic model are the relevant intermediate states and rates of transitions between those connected states. Framed in network language, the states comprise the nodes and the rates define the edges (Figure 1). The reactive flux through the network is determined by the populations (normalized probability) of the nodes and the weights of the edges (transition rates). The relative flux through different reaction pathways can be considered the percentage of single molecules that carry out the process of interest by traveling through a particular series of intermediates (a particular pathway). Once defined, such a kinetic model maps the flux of population changes, such as ion flow in a channel, under any set of conditions, including equilibrium, steady-state, and non-equilibrium relaxation from a given set of starting populations. The MKM procedure can be broken into 9 steps, which will be successively described below in each section, but iteration through these steps is frequently essential (Figure 2).

**Figure 2.**
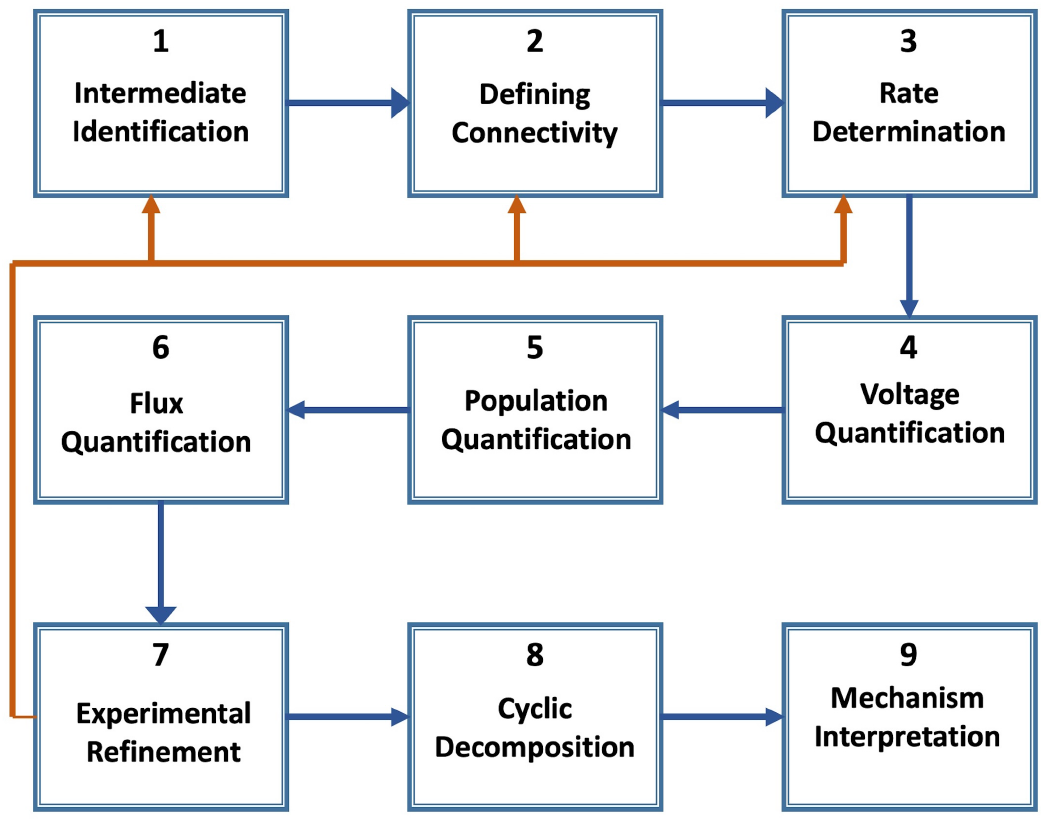
MsRKM process highlighting sequential steps (blue arrows) and iteration (orange arrows).

### 2.1. Intermediate Identification (State Space)

In principle, the selected state space should include all metastable intermediates that are separated from others by rate-influencing barriers, and which contribute above a given threshold value to the ensemble flux for the range of conditions of interest.

This can be a challenging aspect of kinetic modeling since identifying all metastable intermediates requires extensive system characterization—often through a combination of experiments and simulations.^49^ If correctly identified, a kinetic network constructed from these intermediates and their transitions can describe protein behavior under a wide range of experimental conditions. Even when the intermediate state space is not precisely resolved, relevant trends and insights can still be obtained, ideally by comparing several kinetic networks with different state space definitions as demonstrated for the Shaker channel below.

In the models described herein, intermediate states are assigned a binary descriptor (0/1) that is helpful for tracking each feature that can change, such as ion/ligand binding (empty/bound), conformational changes (conformation 1/conformation 2), or chemical state (reactant/product). The number of unique states can of course be increased for features with more than two possible definitions, such as the three competing conformations in a canonical alternating access mechanism (inward-facing, outward-facing, and occluded)^50^. For example, an ion channel with three ion binding sites would be defined as 000 when empty and 111 when bound by three ions. The binary can be reduced to a single number (e.g., 0 for 000 and 7 for 111) for an abbreviated state definition. Retaining the binary description, this yields *N* = 2^*descriptors*^ total states, although some may not be relevant (as demonstrated for ClC-ec1 below). Though not defined in the binary descriptor, MsRKM also allows competing orientations of the protein embedded in a membrane to replicate data from liposomal assays lacking protein orientation. The model assumes a 1:1 ratio of biological to opposite orientations for simplicity. This ratio can be modified for proteins that exhibit preferential membrane orientation or removed altogether for oriented systems such as Shaker.

In this study, the intermediates of ClC-ec1 are defined by three Cl^−^ binding sites (S_out_, S_cen_, and S_in_), two H^+^ loading sites (the lower E203 and upper E148 residues), and the rotation orientation of E148 (up/down) (Figure 3) as detailed in previous studies.^6, 9, 33, 34^ The Shaker channel is modeled with either three or four 4 K^+^ binding sites (Figure 3) located at the S6 ‘gate’ region (S_1_)^39^ and within the selectivity filter at T441(S_2_), G443(S_3_), and G445(S_4_)^38, 51^. All intermediates are included in the Shaker model, but only 48 of the 64 total intermediates are physiologically relevant in ClC-ec1 model since the S_cen_ site cannot be bound by Cl^−^ when E148 is in the ‘down’ conformation^34^.

**Figure 3.**
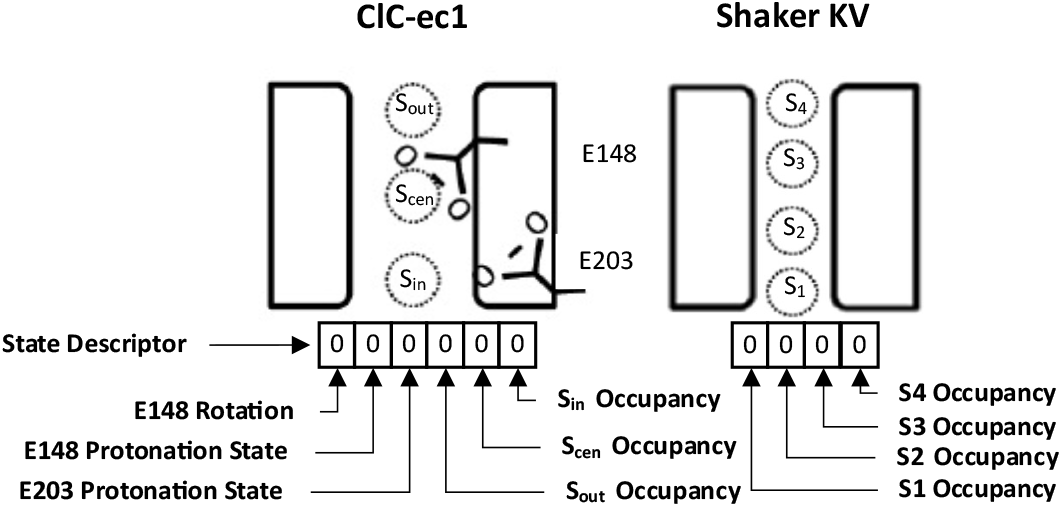
Binary state descriptors for ClC-ec1 (left) and the Shaker Kv channel (right)

It is also possible to simplify a reaction network by grouping states together that exchange more quickly than a user-defined frequency. This can help alleviate numerical challenges and reveal important pathways. Considerable effort has been put into developing robust methods to accomplish this type of network reduction. Since they are not explored herein, the interested reader is directed to spectral clustering^52, 53^, model reduction^54^, and graph theory^55^ for examples of commonly used network reduction approaches.

### 2.2. Defining Connectivity (Transitions)

The reaction network is bidirectional, such that each rare event transition connecting two states/nodes has a forward and reverse directional edge and corresponding forward and reverse rate coefficient. For a fully connected network, this results in N*(N-1) edges and rate coefficients. However, for most systems only certain states will be connected by physically meaningful transitions that contribute to ensemble flux. For example, an ion cannot tunnel through a binding site from Sout to Sint (Figure 3), and transitions involving simultaneous rare-event crossings (e.g., proton uptake to E203 and Cl^−^ release from Sout) are excluded since the lack of coupling between these processes makes the probability of their simultaneous occurrence negligibly small.^34^ In cases where coupling introduces a unique rate for simultaneous transitions, these rates should be included.

For ClC-ec1, this reduces 48 x (48-1) (2256) transitions to 248 transitions. Additionally, some transitions may have degenerate rate coefficients. For example, Cl^−^ uptake to Sout is assumed to be independent of the protonation state of E203 in the presented model of ClC-ec1. This further simplifies the network to 68 unique rate coefficients that describe ClC-ec1. The Shaker model assumes the open conformation of the protein and has no such conformational constraints within the region of the selectivity filter.^56^ However, ions still cannot tunnel (e.g., from S1 to S3) and the current model assumes transitions are not coupled. Future work will explore the role of coupled transitions in Shaker. This reduces the number of transitions from 240 (16*(16-1) transitions from 16 (2^4^) states) to 56 transition rates that were modeled explicitly with no degeneracy assumed between the transitions.

### 2.3. Rate Determination in the Absence of Voltage

Rate coefficients can occasionally be directly measured in experiment^57^, but are most commonly obtained through either bottom-up in silico quantification or top-down refinement based on experimental data. There is a wide array of approaches available to calculate rates from simulations, ranging from quantification of the transition free energy profiles, often called potentials of mean force (PMFs), combined with transition state theory, to the direct tracking of transition probabilities in, for example, Markov state modeling. Often, these methods are accompanied by methods to identify the proper reaction coordinates and to improve the statistical convergence via adaptive or enhanced free energy sampling^58^. The choice of the method depends on the nature of the transition. For example, when chemical reactions are involved, this step can require multiscale treatment, e.g., via quantum mechanics/molecular mechanics (QM/MM)^59^ simulations or reactive MD, such as the multiscale reactive molecular dynamics (MS-RMD),^60^ particularly developed for long-range proton solvation and dynamics. When highly interfacial or ionic systems are involved, polarization effects can be important, requiring polarizable MD. Taking ClC-ec1 as an example, multiple computational methods were used to calculate rates for Cl^−^/H^+^ exchange. MS-RMD was used to characterize the PMF of proton transport (PT) through the central region between E148 and E203,^34^ as well as between E148 and bulk solution^61^. In both cases, the influence of Cl^−^ occupancy at the S_cen_ binding site was explicitly accounted for. All-atom MD with a DRUDE polarizable force field^62, 63^ was used to determine multidimensional PMFs for Cl^−^ transport to capture the influence of ion-ion repulsion.^6^ Once the PMF has been characterized, transition state theory^64^ can then be used to convert the free energy barrier between metastable states and the dynamics of the system in the reactant well into the rate coefficient through the Eyring-Polanyi equation^65^:

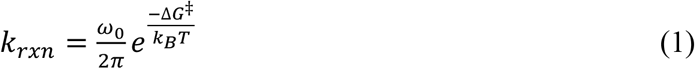

where *k*_*B*_ is Boltzmann’s constant, T is temperature, and Δ*G*^‡^ is the free energy barrier height. The fundamental frequency *ω*_0_ is specific to the transition and quantifies the frequency with which the reactant attempts to cross the transition barrier. It can be approximated from the second derivative of the free energy profile in the reactant well:

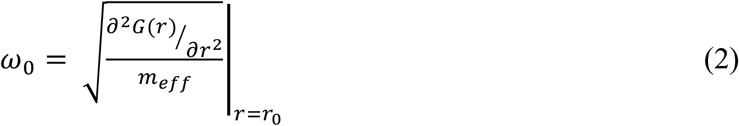

Here 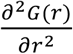 is the second derivative of the free energy G(r) with respect to the reaction coordinater, evaluated at the local minimum in the PMF (*r*_3_). The effective mass (*m*_*eff*_) can be calculated using the thermal velocity of an ion, *m*_*eff*_⟨*v*^2^⟩ = *k*_*B*_*T*/2, where ⟨*v*^2^⟩ is calculated at *r*_0_. While the thermal velocity is not directly dependent on the reaction coordinate, the effective behavior of the ion along the reaction coordinate can vary due to variations in the local environment and potential energy surface. This is particularly important for transitions involving high-energy barriers, where the local thermal environment can influence the rate and mechanism of the reaction. ^66^

Finally, a few rate coefficient values were estimated from alternative methods. BD simulations were performed to estimate Cl^−^ uptake/release rates to/from S_in_ and S_out_.^6^ E148 and E203 protonation and deprotonation rates when S_out_ or S_int_ is occupied^6^ were estimated using the relative change in pK_a_ calculated from PropKa.^67, 68^

Regardless of how they are obtained, these initial rate coefficients act as a crucial ‘seeds’, helping to identify the correct region of parameter space while the error in their estimation can be used to set bounds for parameter optimization. Since biological systems operate in thermodynamic cycles, an additional thermodynamic constraint on the rate coefficients can help ensure good seed values. At chemical equilibrium, the rate constants for a thermodynamic cycle have the following relationship:

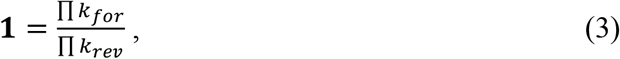

where k_for_ are all rate constants taken in one direction around the cycle and k_rev_ are taken in the opposite direction. Since rate constants do not change as concentrations shift away from equilibrium, this relationship remains valid away from equilibrium as well, assuming no voltage, pH, or conformational changes occur. Using this relationship to precondition values for rate coefficients helps to provide a variety of optimization initiation points within the error of the rate coefficients while guaranteeing that none stray from thermodynamic feasibility.

### 2.4 Voltage Quantification

One important consequence of kinetic selection can be the shifting of mechanistic pathways due to reaction conditions, as demonstrated by molecular motors under high/low ATP/ADP concentrations.^17, 18^ A few candidate proteins^20, 49^ have demonstrated that this also happens outside of the realm of chemomechanical coupling. Most notable for the purposes of this paper, ClC-ec1 has shown that different pH^9^ and Cl^−^ gradients^6^ generate significant mechanistic pathway changes. However, this work did not explicitly factor in the influence of voltage. The transmembrane potential includes both chemical and electrical gradients and the latter must be included to understand the role of kinetic selection in ion channels and transporters. With MsRKM, we seek to capture the influence of the transmembrane electrochemical gradient on the rates of ion transport in a system- and voltage-specific manner. Significant work has gone into developing and validating computational methods to quantify the influence of voltage on substrate transport.^69-72^ We focus herein on methods that combine the calculation of free energy profiles for ion permeation with the dimensionless coupling profile that captures the influence of voltage as a function of the permeation reaction coordinate (often simply the z-axis) across a given channel or transporter of interest.

#### 2.4.1 The constant electric field method

The transmembrane voltage, or electric potential, originates from an asymmetric distribution of ions that is actively maintained by ion pumps and transporters. This charge imbalance creates an electric field that pulls complementary ions to the membrane-solution interface, localizing the voltage drop to the transmembrane and interfacial region. Unfortunately, capturing a charge imbalance in MD simulations that employ periodic boundary conditions (PBC) isn’t possible since the solution on either side of the membrane is continuous.

A commonly implemented solution to this problem is to impose a uniform electric field, E, perpendicular to the membrane plane (conventionally through the z-axis) creating a voltage difference over the length of the simulation.^73-76^ The theoretical foundation of this approach has been well described^77^ and validated^76^. In essence, it mimics experimental ion exchange electrodes holding the solutions on either side of the membrane at different voltages via an electromotive force. The system is conceptually broken down into bath surrounding a subsystem, where the electromotive force in the bath creates a linear “applied potential” and thus constant electric field across the subsystem close to the membrane (i.e., the simulation box). The response of the system, redistributing ions to either side of the membrane, creates the “reaction potential”. The resulting total potential across the box, calculated as a sum of the reaction and applied potentials, concentrates the change in potential across the membrane^76, 77^ (Figure 4).

**Figure 4.**
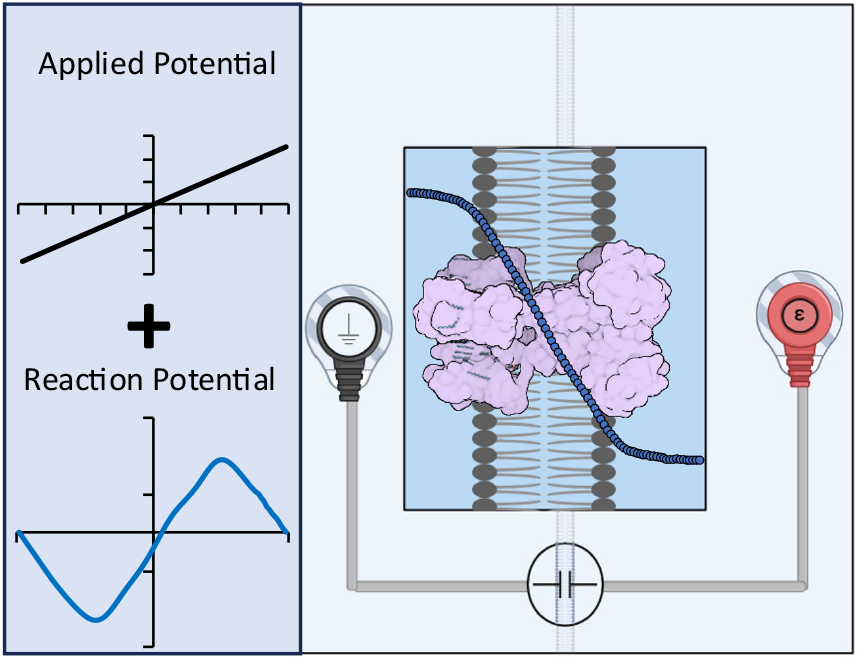
Membrane voltage captured by treating simulation box as a subsystem surrounded by a bath with ion exchange electrodes (right) inducing an applied potential (upper left). The system responds by rearranging charged species to create its own reaction potential (lower left). The summation of the applied and reaction potential localizes the total potential drop to the membrane region (right), mimicking the real membrane voltage created by a charge imbalance localized at the membrane interface.

#### 2.4.2 System-specific voltage

The dimensionless coupling factor φ(z), also called the fraction of the membrane potential, couples the charge movement of the region close to the membrane and the transmembrane voltage applied by the electromotive force in the bulk solution. φ (z) can be determined from multiple methods; some using explicit solvent^77^ and some using the linearized PB-V continuum approximation^78^. The method used in this paper, the linear coupling method^79^, calculates φ(z) by analyzing the ensemble averaged electrostatic potential with PMEpot^75^ module in VMD^80^. This yields a 3-D electrostatic potential map (Δ*ϕ*_*elec*_(*x, y, z*)) which is then integrated over the x and y dimensions to yield Δ*ϕ*_*elec*_(z). φ(z) can then be calculated as the difference between two Δ*ϕ*_*elec*_(z) values:

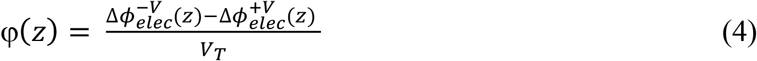

where 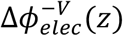 and 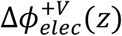 are the ensemble averaged electrostatic potentials for simulations run with a negative and positive applied voltage of equal magnitude; *V*_E_ normalizes the expression^79^. This approximation provides a useful description of the voltage profile along the z-axis of a simulated protein and captures the shift in an ion permeation free energy profile for model systems surprisingly well^76, 79^. However, it cannot describe significant voltage-induced conformational changes for which other methods are more appropriate^77-79^. Alternative approaches to calculating φ(z) and including voltage-induced conformational changes in the rate response will be the focus of future work. For this work, we limit our analysis to conditions under which we do not expect large voltage-induced conformational changes, or equivalently optimize unique solution sets for different conformations.

#### 2.4.3 Obtaining a voltage profile via simulation

Two simulation systems were used to determine φ(z): one that consists of the ClC-ec1 dimer (PBD:1OTS)^25^ with the internally bound Cl^−^ ions removed from each monomer saturated with 163 POPE lipids and another of a protein-less membrane bilayer comprised of 231 POPE lipids used to represent a generic membrane-bound protein in the simple channel system. Both systems were saturated with 300mM NaCl and solvated with TIP3P^81^ water in a 92Å x 92Å x 100Å box with periodic boundary conditions. The CHARMM36^82^ forcefield was used for protein, and ions, while CHARMM36m^83^ was used for the phospholipid membrane. The protonation states of all residues were determined based on previous pKa calculations^84^ on the same crystal structure (E148, E113, and D417 were protonated), while standard protonation states were chosen for all other residues. The influence of pH on ClC-ec1 titration states and the complementary kinetic descriptions will be the focus of other work.

Minimization and equilibration were carried out with NAMD2.14^85^ using a seven-step protocol suggested by CHARMM-GUI^86-88^. Both systems were minimized for 10,000 steps of conjugate gradient energy minimization with restraints on the protein’s heavy atoms (10 kcal mol^-1^ Å^-2^ harmonic positional restraint) without SHAKE^89^. A six-step equilibration was then carried out, gradually removing position restraints on the lipid, protein backbone atoms, and ions. To prevent the spontaneous uptake of Cl^−^, additional wall potentials were placed at each side of a 35 Å x 30 Å rectangular box located at the channel mouth on each monomer, which was large enough to cover the entire area of the pore. The SHAKE algorithm^89^ was introduced to constrain hydrogens in the 2^nd^ equilibration step. The first two steps used the NVT ensemble, with the last four switching to the NPT ensemble. The first three steps were 125 ps, each with a timestep of 1 fs. The final three steps, each lasting for 0.5 ns with a timestep of 2 fs, decreased the force constant for backbone restraints from 1.0 kcal mol^-1^ Å^-2^ to 0.5 kcal mol^-1^ Å^-2^ to no positional restraints. The temperature was maintained at 300 K using a Langevin thermostat with a relaxation time constant of 1 ps in the NVT ensemble. Anisotropic pressure scaling with a Langevin piston barostat was applied to maintain a constant pressure in the NPT simulations, with a damping coefficient of 1 ps. Long-range electrostatics were evaluated using the particle-mesh Ewald method^90^. The cutoff of the van der Waals and short-range electrostatic interactions was set to 12 Å which was smoothed with switching functions.

Each system (ClC-ec1 and the membrane only) next underwent two separate voltage equilibrations using a constant electric field (equal to a +500 mV and –500 mV voltage drop across the membrane) applied perpendicular to the membrane along the z-axis. The voltage equilibrations employed the same NVT ensemble as described above. Harmonic restraints with a spring constant of 2 kcal mol^−1^ Å^−2^ were applied to protein backbone atoms to prevent large conformational changes. Each system underwent 10 ns of simulation, with the last 2 ns used for analysis. Snapshots taken every 2 ps were analyzed in the VMD^80^ PMEpot plugin^75^ to compute *ϕ*_*elec*_(*x, y, z*), and φ(z) was calculated using equation 4 (See figure 5).

**Figure 5.**
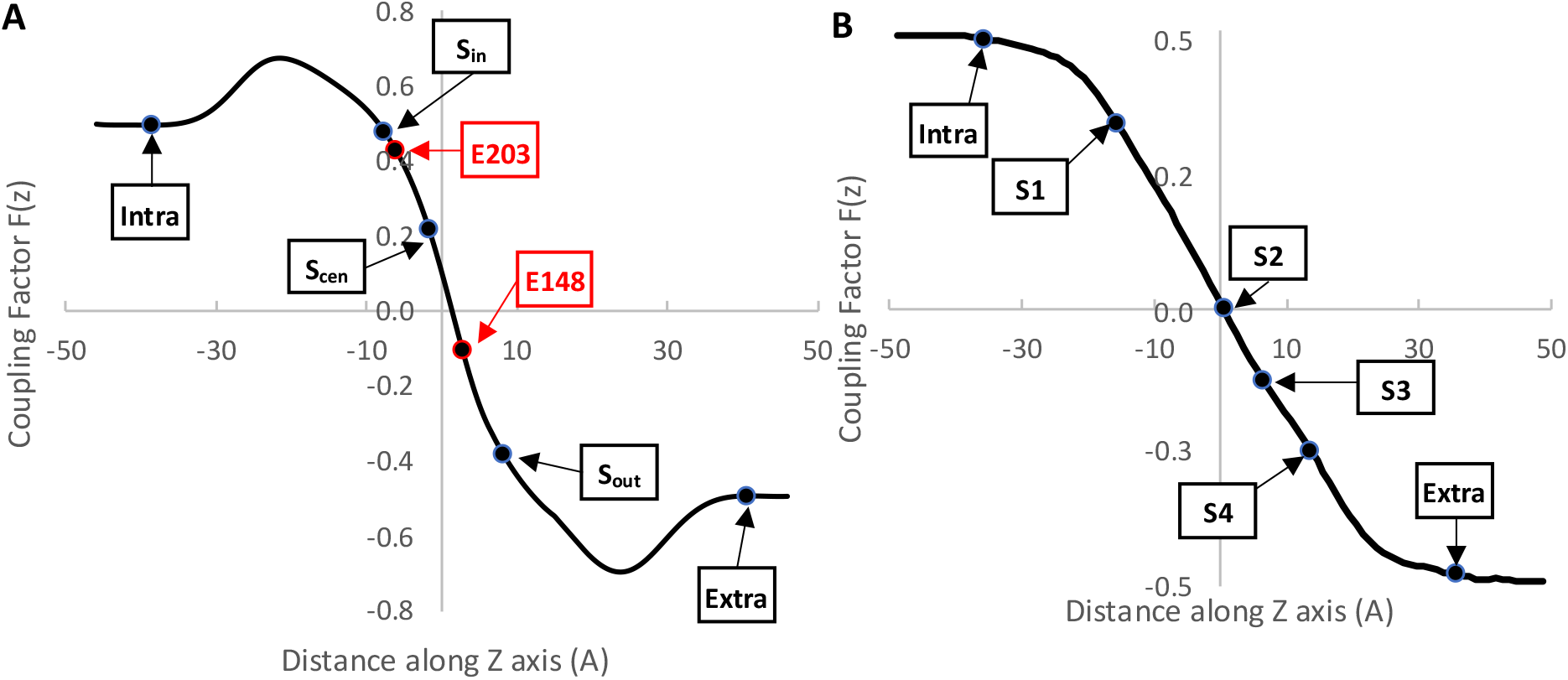
Voltage coupling factor derived from PMEpot from simulations with ClC-ec1 present (A) and absent (B). Important site locations for ClC-ec1 (A) and Shaker (B) have been mapped on the coupling factor used for each model.

#### 2.4.4 Voltage-responsive rates (Derivation)

An alternative way to derive a dimensionless voltage coupling profile, φ(z), is to take the difference between ion translocation PMFs run at different voltages. This so-called *W-route*, which has been detailed in previous studies^77, 78^, can be used to describe an ion transport PMF under the influence of a transmembrane potential as:

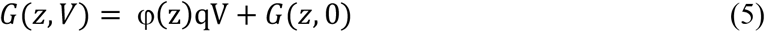

Rates of rare-event transitions are defined by the difference in free energy between meta-stable intermediates and transition barriers (ΔG^‡^). Using equation 5, we can estimate the voltage-induced changes in ΔG^‡^ using the relationship:

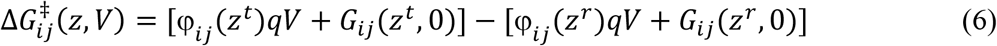

where 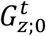 and 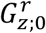 represent the free energy at the transition barrier and reactant basin,respectively. Placing equation 6 into equation 1, we can derive a relationship between the rate coefficients under different transmembrane potentials:

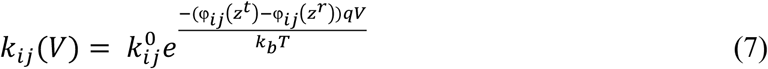

where 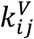 and 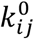 represent the rate coefficients for a transition from state i to j with and without the influence of voltage and k_b_ is in units of eV K^-1^.

### 2.5. Population Quantification

Once states and transitions are defined and rate constants are obtained, the time-dependent state populations can be quantified with a kinetic master equation:

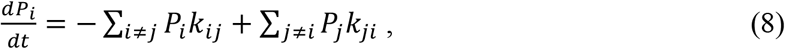

where *P*_*i*_ is the population of state *i*, and the summation includes all other states *j* connected via a transition to state *i*. This set of linear differential equations can be cast in matrix form:

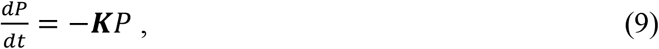

where **K**, the rate matrix, has negative off-diagonal components **K**_ji_ = -*k*_*ji*_ and positive diagonal components **K**_ii_ = Σ_*j*≠*i*_ *k*_*ji*_ for all states *j* connected to state *i* and **K**_ji_ = 0 otherwise. **K** describes the physical flow or motion of ions and molecules in continuous time. **K** is directly related to **T**, the transition matrix used in Markov State Models (MSM). **T** describes the abstract flow of probability between states in discrete time and is the matrix exponent of **K**:

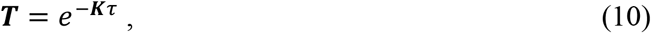

where τ is the timestep size used in the MSM.

By setting *dP/dt* = 0, the system can be solved for the equilibrium population of states. This equilibrium solution is required to be microscopically reversible with detailed balance such that P_i_ k_ij_ = P_j_ k_ji_. In biological systems, where ion/substrate concentrations are often maintained physiologically, the system may not reach equilibrium due to constant influx to and efflux from the local environment surrounding the protein of interest. In this situation, the system may instead approach a non-equilibrium steady state with constant populations and flux:

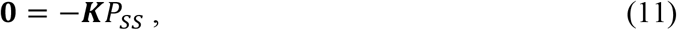

where 0 is a zero column vector with rows equal to the rank of the rate matrix and *P*_*ss*_ is the column vector containing the steady state populations. Steady-state solutions do not obey detailed balance. Adding these sources and sinks into the system of equations reduces the number of independent equations, necessitating the replacement of one state’s rate equation with a normalization condition, and the equivalent row in the zero vector with the desired norm. The steady-state solution corresponds to the solution of an eigenvalue-eigenvector problem with an eigenvalue of zero, the corresponding normalized eigenvector providing the steady-state populations. Other eigenvalues-eigenvector pairs correspond to the dynamics of the system as it relaxes to equilibrium or steady-state, with large eigenvalues corresponding to slower relaxation modes.

The kinetic master equation also permits general time-dependent solution, allowing the evolution of the populations in time to be observed. If the environmental conditions are permitted to vary, these solutions show populations evolving toward equilibrium. If they are held constant, the populations will instead evolve toward a non-equilibrium steady state.

### 2.6 Flux Quantification

The net number of ions transferred through any transition is computed as:

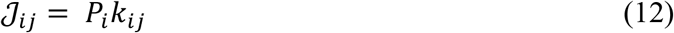

Where *𝒥*_*ij*_ represents the flow in 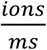 going from state i to j. The net flow going through the protein is determined by summing all transitions to/from the inner-most or outer-most ion binding sites as:

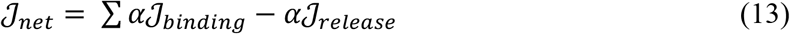

where *α* defines the directionality of the transitions as either outflux (*α* = -1) or influx(*α*= 1). For example, when analyzing Cl^−^ influx of ClC-ec1, equation 13 becomes:

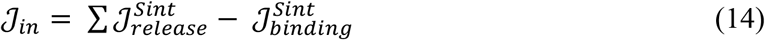

where we have defined negative net flux to be ions flowing out of the cell and positive net flux as ions flowing into the cell.

### 2.7 Experimental Refinement

#### 2.7.1 Limitations of bottom-up kinetic modeling

Although the rigor and accuracy of simulation-based thermodynamic and kinetic analyses are impressive and rapidly improving, systematic^91^ and statistical^92^ error is inevitable. In principle, obtaining a well-behaved kinetic model from bottom up is possible and has been demonstrated for select systems^28^. Frequently, however, compounding errors^93-96^ will quickly throw off a kinetic solution. The same challenge has been observed in microkinetic modeling, an arguably more advanced field focused on modeling chemical reactions and heterogeneous catalysis^97, 98^. To remedy this, a top-down quantification of the kinetic parameters by optimization with experimental results is performed with optimization boundaries set to the calculated error of MD simulations.

#### 2.7.2 Integrating experimental data

The solution space of kinetic networks tends to be vast, with the number of unknowns (rate coefficients) typically outnumbering the number of known data points with which to refine them. The ability to refine a solution thus hinges on two things: 1) good initial rate coefficients (seeds) with well-defined boundary conditions, and 2) the amount and quality of data that can be used as optimization constraints. High quality experimental data is not only that with small error bars, but also data that uniquely constrains the optimization. For example, five sets of IV curves under similar conditions will not refine the solution as well as three IV curves under highly different conditions and two liposomal flux assays. In general, adding additional experimental data will further reduce the solution space and improve the solution.^49^ An overdetermined system tends not to be a problem when extracting generalizable trends if the results are consistent. In such a case, some equations running within the optimization may end up superfluous, resulting in poor optimization performance. If the overdetermined model produces inconsistencies or a lack of solution, overfitting can be handled with regression techniques to reduce the total number of constraints or equations.

However, it is important to note that it is possible to get optimization solutions that show different protein behaviors or a lack of optimization solutions regardless of the system type. In our experience, this has often been due to some experimental inconsistencies or clashing physical realities with other experimental results placed in the optimization. For example, the presence of an undetected ion leak could lead to an I-V curve that appears similar in profile to other I-V curves but has designated a reversal potential where an efflux assay, under identical experimental conditions, has non-zero flux. Difficulty finding a solution may also be due to assumptions made in the model-building process, especially if one is reducing to the minimum possible states to describe a network^99^.

#### 2.7.3 Loss function design

The choice of optimization methodology is critical to the success of the optimization. To allow different methodologies in the fitting of experimental data, SpotPy^100^ global, SciPy^101^ global and local and user-defined optimization algorithms can be used. Though not the focus of this study, extensive work has been done to examine the best optimization algorithms for biological-based networks.^102-104^ The selected optimization method changes the intrinsic rate coefficient values while minimizing a least squares loss function incorporating experimental data. Each experimental data point reflects different pH and ion concentrations for the intra-and extracellular sides of the membrane and different values of ΔΨ that produce an experimental observable to be compared to. We currently support several experimental observables in the loss function, such as liposomal flux assays, I-V curves, and orientation-based blockage studies. Optimizations based upon flux through the liposomal system also allow for the optimization of net flux through different orientations separately, i.e., net flux through the biological orientation only. The optimization also allows for flux component differentiation to be accounted for within the loss function, such as differential orientation influx/outflux, total influx (defined as all flow moving inwards from both orientations), total outflux (defined as all flow moving outwards), and the ability to turn the microscopic reversibility term on or off for functionally irreversible systems. The model also allows for the user to optimize to normalized flux values such as normalized IV curves, normalized IV curves with differential orientation preferences, or normalized flow assays. Microscopic reversibility has been represented as a mean squared error function normalized over the total number of transitions within the loss function:

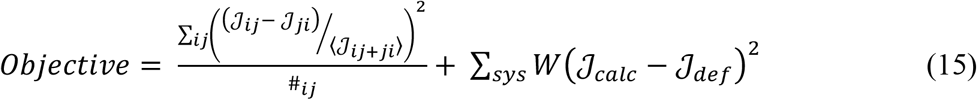

where microscopic reversibility is included in the first term of the objective function, the second term calculates the least squares difference between the flow values calculated through MsRKM and the user-defined values for every experimental system input. MsRKM also allows for the weighting of different experimental values within the loss function dependent upon the experimental type represented by W in the equation above.

### 2.8 Cyclic Pathway Decomposition

Once a kinetic solution is obtained, the relative contribution of different pathways must be quantified. This is a nontrivial task as the flux into and out of any given node can contribute to multiple pathways. Although it is more straightforward to identify the dominant path by tracking the largest minimal flux transition,^105^ this does not quantify the relative contributions of all pathways. To accomplish this goal, we previously developed CycFlowDec^106^, an algorithm that quantifies the expected flux through simple cycles (each containing any given node no more than once) in a closed network. Based on established methods in Markov circulation theory,^107, 108^ the algorithm in its simplest form decomposes the total flux into relative cycle contributions based on random walks generated from Markov transition probabilities. It is then made efficient for complex networks by using a percolation with burn-in and minimum contribution tolerance. Herein we implemented CycFlowDec^106^ for ClC-ec1 and Shaker with a 100,000-step percolating algorithm with a burn-in of 4500 steps, a flow tolerance of 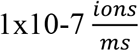, and an MRE cutoff of 9×10^−3^.

### 2.9 Mechanism Interpretation

The above analyses provide a full network description of the system’s flux through various intermediates, which can be tracked under the equilibrium, steady state, or non-equilibrium conditions for which the rates remain applicable. Thus, the mechanism can be compared under varying reaction conditions and the role of competing pathways can be delineated. Understanding the mechanism at the molecular level, and hence the structure-dynamics-function relationships that control the process efficiency and selection between competing pathways, is of course benefitted by simulations probing the transition ensembles. Simulations are also best suited to probe the impacts of point mutations on rate-influencing transitions. In addition, MsRKM can be used to probe and compare the network descriptions that differentiate wild type from mutant behavior, as demonstrated for Shaker below.

#### 2.9.1 Pathway diagram visualization

Network flux can be tracked graphically as a function of time, or via network diagrams for equilibrium or steady-state conditions. For the latter, all significant pathways of a solution set can be plotted in a corresponding full network diagram or a subset of dominant or related pathways can be plotted in a pathway diagram. In this work, each diagram uses a series of cartoon representations of either ClC-ec1 or the Shaker channel states connected by arrows that represent transitions. The depictions include (Figure 6):

**Figure 6.**
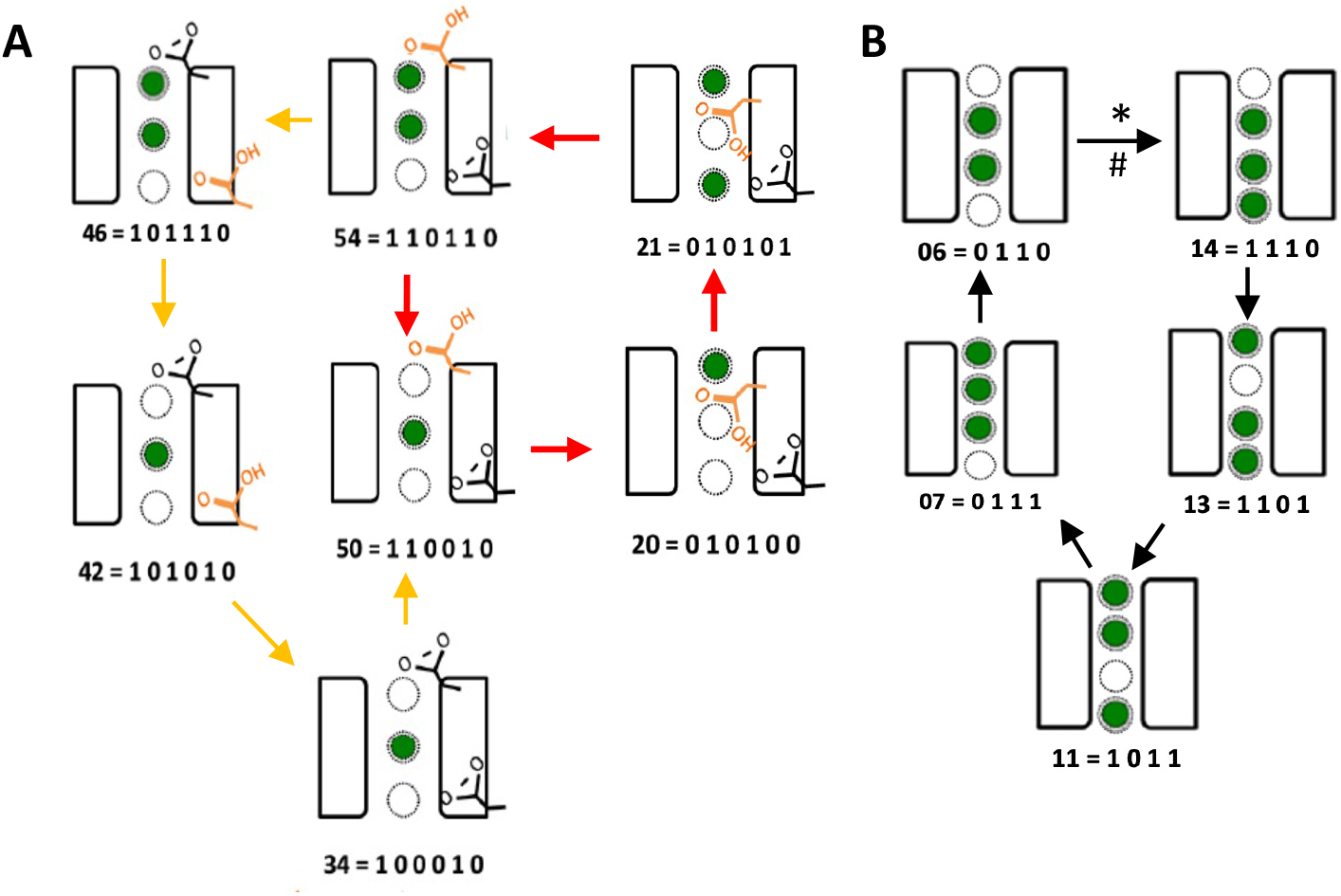
Example Cycles for (A) ClC-ec1 and (B) Shaker

1. the non-interacting portion of the protein represented by two empty rectangles,
2. three dotted circles representing the ion binding sites that are filled 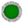 when occupied or empty 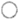 when unoccupied,
3. H^+^ binding sites for ClC-ec1, E148 and E203 either protonated 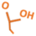 or deprotonated 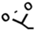,
4. E148 rotating “up” 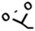 towards the extracellular side or “down” 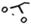 towards the central Cl^−^ binding site (S_cen_) of ClC-ec1,
5. the ‘Biological’ orientation of the protein with the extracellular solution on top and intracellular solution on bottom and vice versa for the “Opposite” orientation,
6. Flux Limiting Steps (FLS) of the cycle denoted with * and Rate Limiting Steps (RLS) denoted with #.

## 3. RESULTS AND DISCUSSION

### 3.1 Model Robustness and Predictions

A central challenge in kinetic modeling is narrowing the solution space. Typically, the problem is underdetermined with far more parameters (rate coefficients) than known data to fit. Thus, kinetic modeling hinges on the quantity and quality of data available. For systems with a range of experimental results (often top-down macroscopic data, but sometimes rates for specific transitions) and simulation-derived kinetic parameters (bottom-up molecular data), MsRKM can display robust predictive power—not only narrowing the solution space but also demonstrating when the underlying network (state space definition) is insufficient (as demonstrated below).

ClC-ec1 as a case study demonstrates the need for experimental and simulation-based data to determine robust solutions. For this system, previous studies employing state-of-the-art multiscale and polarizable simulations quantified most of the ion transition rates with reported errors in the corresponding free energy barriers ranging from 0.8 to 2 kcal/mol^6, 33, 34^. The remaining rates were estimated based on pKa calculations. Despite significant effort to quantify these rates as accurately as possible, they produced unphysical results when placed into a kinetic model due to compounding errors. Note that even small errors in free energy profiles lead to larger errors when exponentiated to estimate rates. Additionally, new work has pointed to a significant conformational change at low pH values that was not accounted for in the simulations^29^. During solution refinement, MsRKM accounts for these errors by adjusting rate coefficients within user-defined boundaries of their reported error to produce a kinetic model consistent with macroscopic and molecular data. These adjustments can be essential, both to meet the requirement of microscopic reversibility and to fit experimental data. Interestingly, even before the conformational change had been published, the solutions that were able to fit the data at low pH absolutely required opening up the error bounds on several rates in a manner that is consistent with the low pH structure. At the time, it was not clear why. The details of these models and their implications will be the focus of separate work. However, the process of determining these models demonstrates how integrating data from both simulations and experiment can be essential to identify robust kinetic solutions.

Starting from the simulation-based seed rates, the solution refinement for ClC-ec1 included constraints of: 1) microscopic reversibility at 0mV with pH 3:3 and Cl^−^ 300:300; 2) five current values at pH 4:4 and a Cl^−^ gradient of 30:300 (Figure 7B, red IV curve^109^) and at pH 5:5 with a Cl^−^ gradient of 40:300 (Figure 7D, light blue IV curve)^31^; 3) five efflux assay values at pH 4, 4.5, and 5^109^; 4) five influx assay values at pH 4, 4.5, and 5^109^; and 5) only two specific reversal potential^26^ values—one at symmetric pH 5:5 and a Cl^−^ gradient of 45mM:300mM (from Figure 7C, top) and the other at pH 5:3 with symmetric Cl^−^ concentrations 300mM:300mM (from Figure 7C, bottom). This combination of experimental data limited the solution space significantly, resulting in an MsRKM model that then successfully predicts: 1) the remaining normalized I-V curves under different conditions (Figure 7B, blue, green, and yellow IV curves)^30^, 2) the remaining reversal potential values for concentration gradients evaluated at pH 3 and 5 (Figure 7C Top), and 3) pH gradients at symmetric 300mM Cl^−^ concentration (Figure 7C Bottom). Since these assays are different from the assays included in the objective function, this offers the first essential validation check for solution robustness.

**Figure 7.**
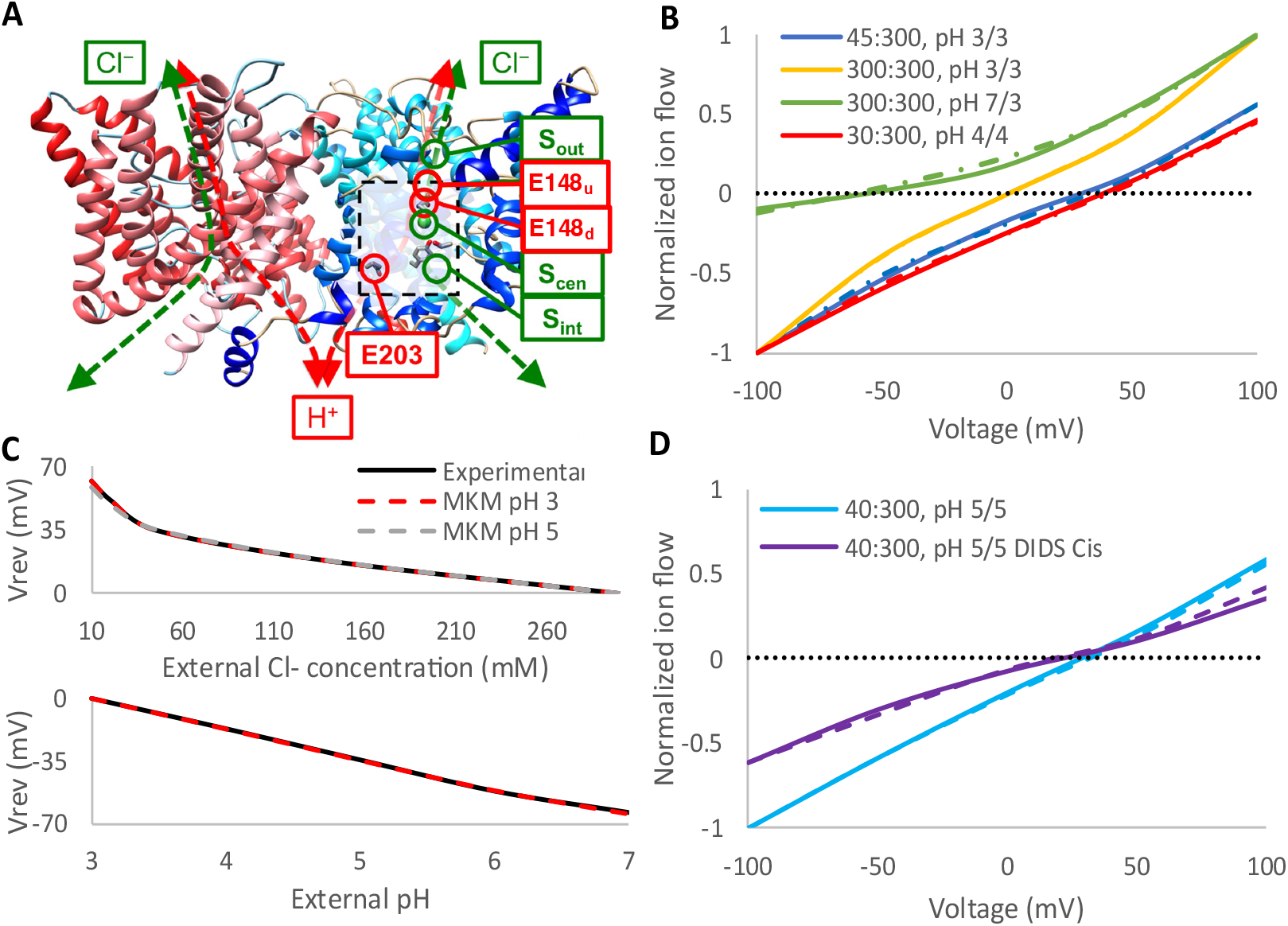
(A) ClC-ec1 highlighting ion binding sites. (B-D) MsRKM results (dashed lines) compared to experimental data (solid lines) for ClC-ec1, highlighting the model’s ability to replicate IV curves (B,D) and predict reversal potentials (C) over a range of conditions. Experimental ion concentrations are reported as extracellular: intracellular with Cl^−^ in mM and H^+^ in pH. In the reversal potential assays (C), either extracellular Cl^−^ was varied with intracellular Cl^−^ at 300 mM, maintaining symmetrical pH (Top), or extracellular pH was varied with internal pH 3, maintaining symmetric 300mM Cl^−^ concentrations (Bottom). (D) MsRKM prediction of directed current (purple) through one-orientation via DIDS-based inhibition at the intracellular side of ClC-ec1.

To further benchmark MsRKM solutions, it is important to test their ability to replicate experimental results for assays that are increasingly different. As a word of caution, the model should only be extended within the conditions under which the rates and rate responses are valid (see Methods). Thus, to further test the ClC-ec1 solutions we modeled a blockage assay^31^ in which stilbenedisulfonate 4,4-diisothiocyanatostilbene-2,2′-disulfonic acid (DIDS) incompletely blocks the internal side of ClC-ec1, providing the only reported estimate of directional flux through the transporter. The opposite orientation is unaffected, so 100% of opposite flow is included, while the biological side is incompletely blocked, so 6% of the biological flow is included. With this single correction to the flows, MsRKM replicated the DIDS-blockage assay normalized I-V curves (Figure 7D). This is a striking result. Although further model testing is always warranted, the ability to fit a weighted unidirectional flow supports the model’s validity and ability to predict ClC-ec1 current under the broad range of conditions tested. This example demonstrates the potential robust and predictive power of MsRKM when the solution space is sufficiently refined and the response of the rates to the range of conditions modeled is properly included.

### 3.2 Top-Down Modeling with MsRKM

Even when specific rate constants are not known, MsRKM can identify robust and predictive models as long as there is sufficient molecular insight to define the state space and experimental data to narrow the solution space. Although the most insight is gained by combining top-down modeling from experimental data and bottom-up characterization of the dynamics and rates of individual transitions, purely top-down modeling can still generate new biological insight, reveal mechanistic control, and inform experimental design. It additionally provides a framework to fold in increasing bottom-up characterization and test contrasting predictions.

We demonstrate the process of top-down modeling with the Shaker voltage-activated potassium channel (Figure 8A), modeling only the open conformation.

**Figure 8.**
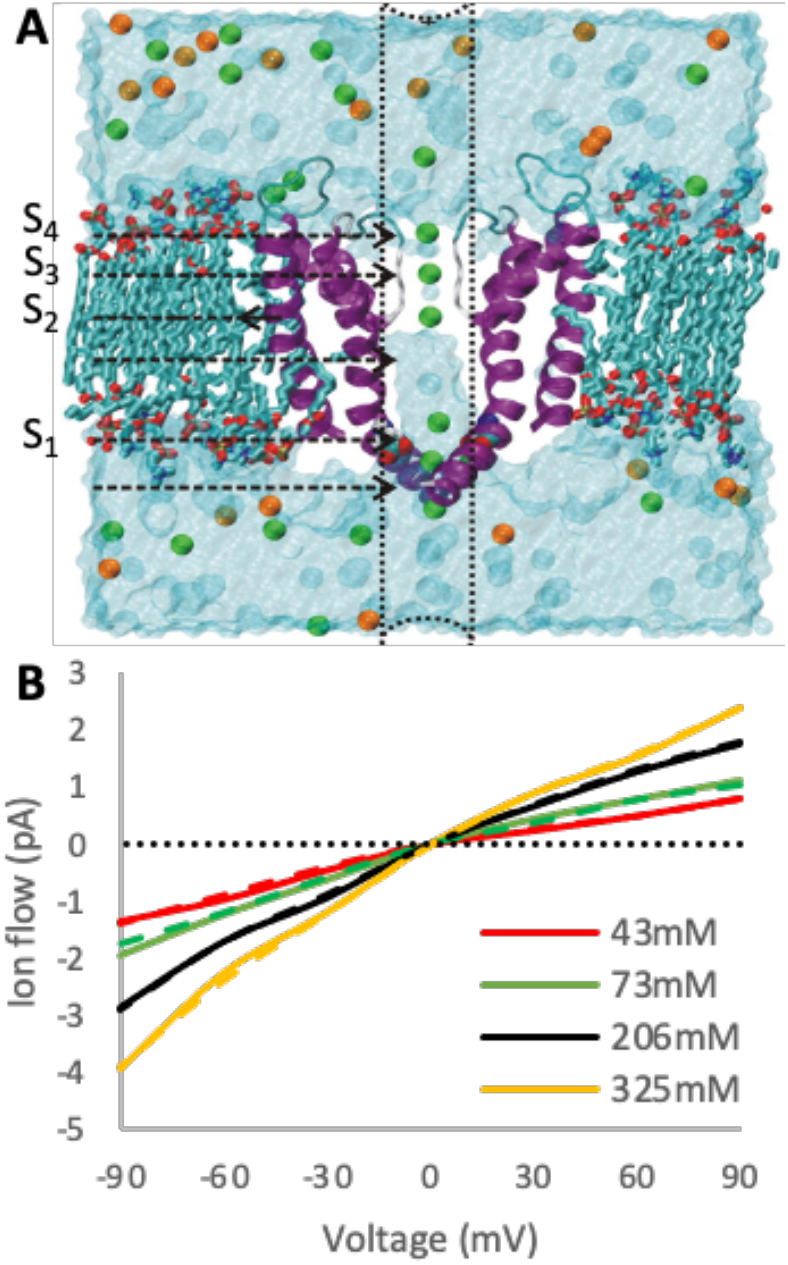
(A) Shaker channel showing site locations modeled in the four-site model (adapted from ^2^). (B) Single-channel I-V curves for the Shaker channel with symmetric K^+^ concentrations, comparing experimental data^1^ (solid lines) to four-site MsRKM results (dashed lines).

We first constructed a four-binding site model of Shaker with one site located at the internal pore entrance (S1) and three binding sites in the region of the selectivity filter, (Figure 5B and 8A) consistent with MD generated K^+^ density plots^2^. We then generated 50 starting rate coefficient sets using a random-number generator to enable the optimizer to broadly sample the solution space. Only K^+^ uptake rate coefficients were constrained with an upper boundary of 1×10^8^ M^-1^s^-1^ (1×10^5^ mM^-1^ms^-1^), approximating the diffusion limit. Sets of rate coefficients were considered successful solutions if they deviated by less than 0.3 pA from experimental single-channel I-V curves (Figure 8B). This criterion was met by 52% (26/50) of optimization runs. However, the was considerable heterogeneity in the resulting 26 Shaker solutions, including different voltage and chemical gradient current profiles, demonstrating the need for additional refinement.

One strategy for solution refinement is to use biophysical insight. Since all K^+^ channels share a strictly conserved selectivity filter but have a wide range in current (from 1.5pA for the Shaker KV channel^1^ to 25 pA in the BK Channel^110^), it has been suggested that the RLS for slower K^+^ channels like Shaker must be outside of the selectivity filter^2^. Specifically, the RLS should be uptake to S1 or transfer from the vestibule into the selectivity filter (S1 to S2). Although it seems unlikely for Shaker, release from S4 could also be considered if structural elements beyond the selectivity filter contribute to the barrier for K^+^ release. Based on these considerations, 13 of the 26 Shaker solutions could potentially be ruled out since their dominant pathways have a RLS in the selectivity filter. The remaining 13 solutions, however, remain viable solutions—11 with uptake to S1 as their RLS, 1 with S1-S2 transfer, and 1 with release from S4.

An alternative strategy for solution refinement is to test the solutions on new experimental data. Following this path, we further refined our solution space by modeling the Shaker-P475D mutation^2^. This mutant is particularly informative because the Pro to Asp mutation lies just inside the inner vestibule near the activation gate and significantly speeds up ion flux, promoting a 6 to 8-fold higher single-channel current. We modeled the P475D mutation using the rate coefficient sets of all 26 possible solutions as seeds in a secondary optimization fitting P475D’s single-channel I-V curves (Figure 9A). Since the mutation is 17 Å below the selectivity filter, we only allowed rate coefficients near the mutation to change between the wildtype (WT) seeds and mutant. This included uptake to S1, S1 release to bulk, and S1→ S2 transfer. Notably, only 3 solutions were able to fit the mutant I-V curves within the <0.3pA acceptance criterion (Figure 9A dashed lines). As expected, the WT RLS of these three solutions is outside the selectivity filter (two are uptake to S1 and one is S1 to S2 transfer at 100 mM Cl^−^). As a less stringent alternative test, we performed a secondary optimization allowing all rates to vary to fit the mutant data. This found an additional 7 solutions. The WT RLS of each of these seven additional solutions remains outside of the selectivity filter. Thus, both the WT and mutant results support the notion that the RLS for Shaker is either ion uptake or transfer into the selectivity filter.

**Figure 9:**
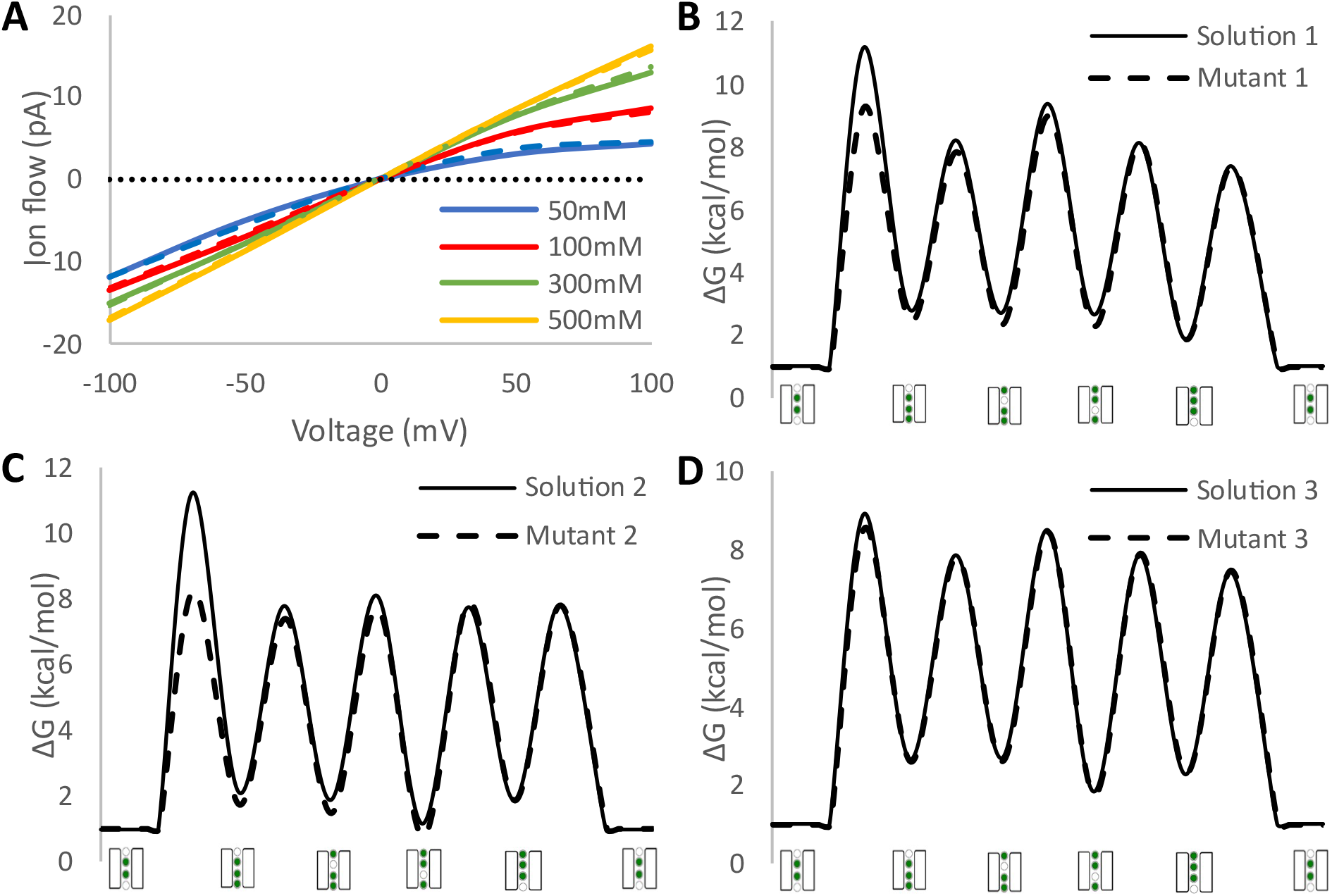
(A) Extension of four-site Shaker models to the P475D mutant with single-channel I-V curves at symmetric K^+^ concentrations comparing experimental data^2^ (solid lines) and MsRKM results (dashed lines). (B-D) Free energy profiles for the dominant pathway in the three best Shaker solutions.

We analyzed their dominant mechanistic pathways at 1:1 mM symmetric K^+^ equilibrium conditions (Figure 9B, C, and D) to differentiate between the three solutions. Analyzing pathways in the outward direction, solutions 1, 2, and 3 all have a RLS of uptake to S1 at this low concentration. Since uptake is rate limiting, the RLSs of the solutions are sensitive to the internal concentration of K^+^. Solution 1 switches its RLS at 400mM from S1 uptake to S2 → S3 transfer. Solutions 2 and 3 switch their RLSs from S1 uptake to S1 → S2 transfer at 496mM and 25mM, respectively. The mutant solutions still have RLSs of S1 uptake, but they reduce the uptake barrier heights by ∼1.8, ∼3, and ∼0.4 kcal/mol for solutions 1, 2 and 3, respectively. Thus, the RLS moves at much lower concentrations. The RLS of solution 1 RLS shifts to S2 → S3 at 9mM, while the RLSs of solutions 2 and 3 shift to S1 → S2 at 2mM and 23mM, respectively. Collectively, the three solutions are unified in RLS at low K^+^ concentration, but they diverge significantly both as K^+^ concentrations increase and as they fit the mutant.

We turned to additional experimental data to further delineate which of the three solutions is best. Like other K^+^ channels, Shaker exhibits saturation of the channel, where increased concentration does not result in increased conduction. Saturation of the channel occurs between 325mM and 600mM, with concentrations >600mM minimally increasing flux through the transporter^1^.We tested the prediction capabilities of the three remaining solutions using the WT saturated I-V curves (Figure 10). All solutions correctly predict the 600mM single-channel I-V curve, but only solution 2 exhibits saturation at 1.15M.

**Figure 10:**
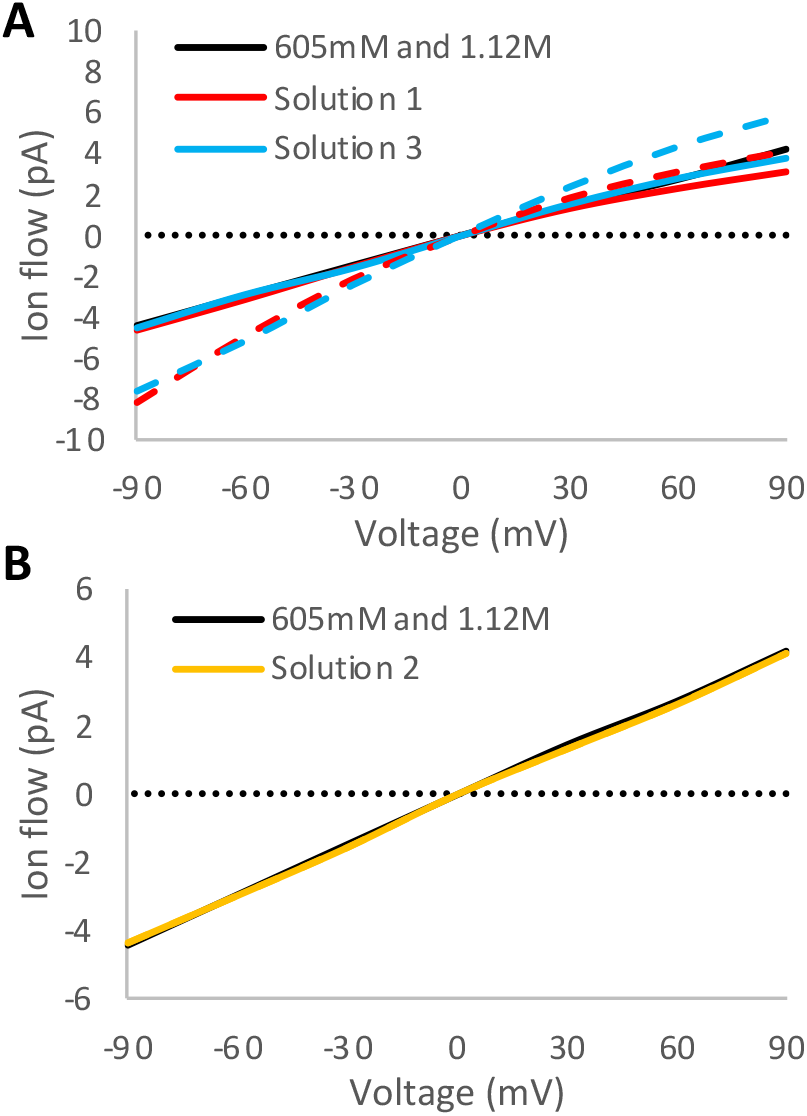
MsRKM solutions compared to experimental data at higher symmetric K^+^ concentrations. (A) Solutions 1 and 3 show clear deviations between 605 mM (solid) and 1.12 M (dashed) results, contrary experiment and (B) solution 2, which are nearly overlapping.

The aforementioned process demonstrates how gradually integrating experimental insights and observations can narrow the solution space, enabling us to identify a single set of rate coefficients (solution 2) that is consistent with experimental data and biophysical insight. However, we caution against assuming additional solutions do not exist. The global optimization procedure explores the phase space stochastically, with a deterministic local optimization to provide locally optimal solutions, but it provides no guarantee that we have found the true global optimum. As solution 2 best describes all experimental data to date, we focus our mechanistic analysis below on solution 2.

### 3.3 MsRKM Can Identify Mechanisms Dominated by a Single Pathway

All previously published MKM models displayed mechanisms with multiple pathways contributing to the net flux^6, 9^ that shifted under different reaction conditions. Thus, a critical test of the MsRKM methodology is not in its ability to identify competing pathways, which it is designed to do, but rather in its ability to eliminate superfluous mechanisms in single-pathway dominant systems. The Shaker four-site model is a good test in this sense. Most of the solutions identified displayed some degree of multiple contributing pathways. However, solution 2 is almost entirely single-pathway dominant, with >99% of ion flow at physiologically and experimentally relevant voltages and concentrations going through the pathway shown in Figure 11A.

**Figure 11:**
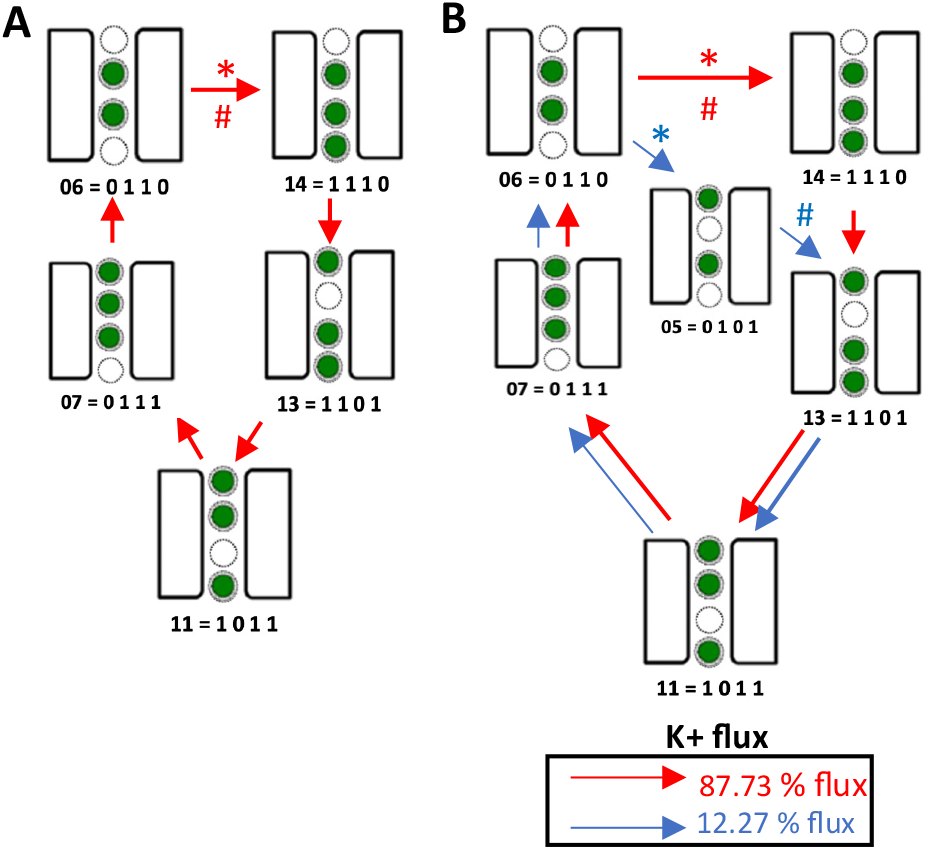
Dominant pathways from solution 2 for WT (A) and mutant (B) at physiological conditions (−25mV, [6mM] ext: [158mM] int). # RLS, * FLS

Interestingly, this mechanism is consistent with that proposed by previously through double membrane computational electrophysiology simulations.^70, 71, 111^ Flux through other pathways only becomes significant at high voltages or concentrations. Although additional mechanistic pathways appear at high concentrations, we observe no significant increase in total flux through the protein at concentrations above 500mM.

In contrast, solution 2 for the P475D mutant does not retain this mechanistic homogeneity (Figure 11B). The secondary pathway of the mutant solution contributes 12.27% to the net flux at -25mV, within normal physiological conditions (Figure 6B blue). This system also shows more voltage-dependent kinetic selection, with the secondary pathway’s contribution to the net flux increasing and decreasing with increasing and decreasing voltage, respectively. Thus, MsRKM correctly identified both a single-pathway dominant mechanism and a multi-pathway mechanism in closely related systems, suggesting that MsRKM is sufficiently flexible to capture the nuances of real data.

While the mechanistic differences between this study’s WT and mutant models are enlightening in exploring MsRKM capabilities, we caution against drawing Shaker-specific mechanistic conclusions. We constructed a simplified Shaker model to find the minimum number of states necessary to replicate experimental data. Most MD and experimentally derived models show additional binding sites^43, 45, 47, 111^ that likely need to be included to draw physiological conclusions. We also used a model membrane voltage coupling factor for Shaker instead of a system-specific profile, such as that shown for ClC-ec1 (Figure 5A). A more detailed MsRKM model of Shaker will be the goal of future work, including more sites, the role of water, and coupled ion transitions. However, if the results described above for our simple four-site model hold in more detailed models, it suggests that Shaker evolved a kinetic network that supports a robust, single-pathway dominant mechanism for K^+^ transport in the ‘open’ configuration.

### 3.4 Model Refinement Provides Clear Feedback between Acceptable Simplification and Oversimplification

While the four-site MsRKM model was capable of fitting all Shaker experimental data to date, theoretical models using fewer sites have provided valuable insights. For example, Mosoco et al. used three potassium ion binding sites in their Eyring model, an early kinetic model that altered barrier heights and well stabilities of a mechanistic pathway,^64^ to explain the P475D mutant’s increased conductance^2^. This suggests that further simplification from the four-site MsRKM could produce viable solutions. To test this, we constructed a three-site Shaker model that combines S2 and S3 (Figure 8A) to form a single site. We then fit the WT Shaker single-channel I-V curves with the same procedure, boundaries, and acceptance criterion detailed above. Out of 100 optimizations, 62 (62%) resulted in a solution that met the acceptance criteria. The I-V curves for one such solution is shown in Figure 12A. These 62 WT solutions were then used as seeds for a secondary optimization to the mutant data, as detailed above. Surprisingly, this secondary optimization did not produce a single viable solution, with the best fit having a 2.96pA average deviation from experimental data (Figure 12B).

**Figure 12.**
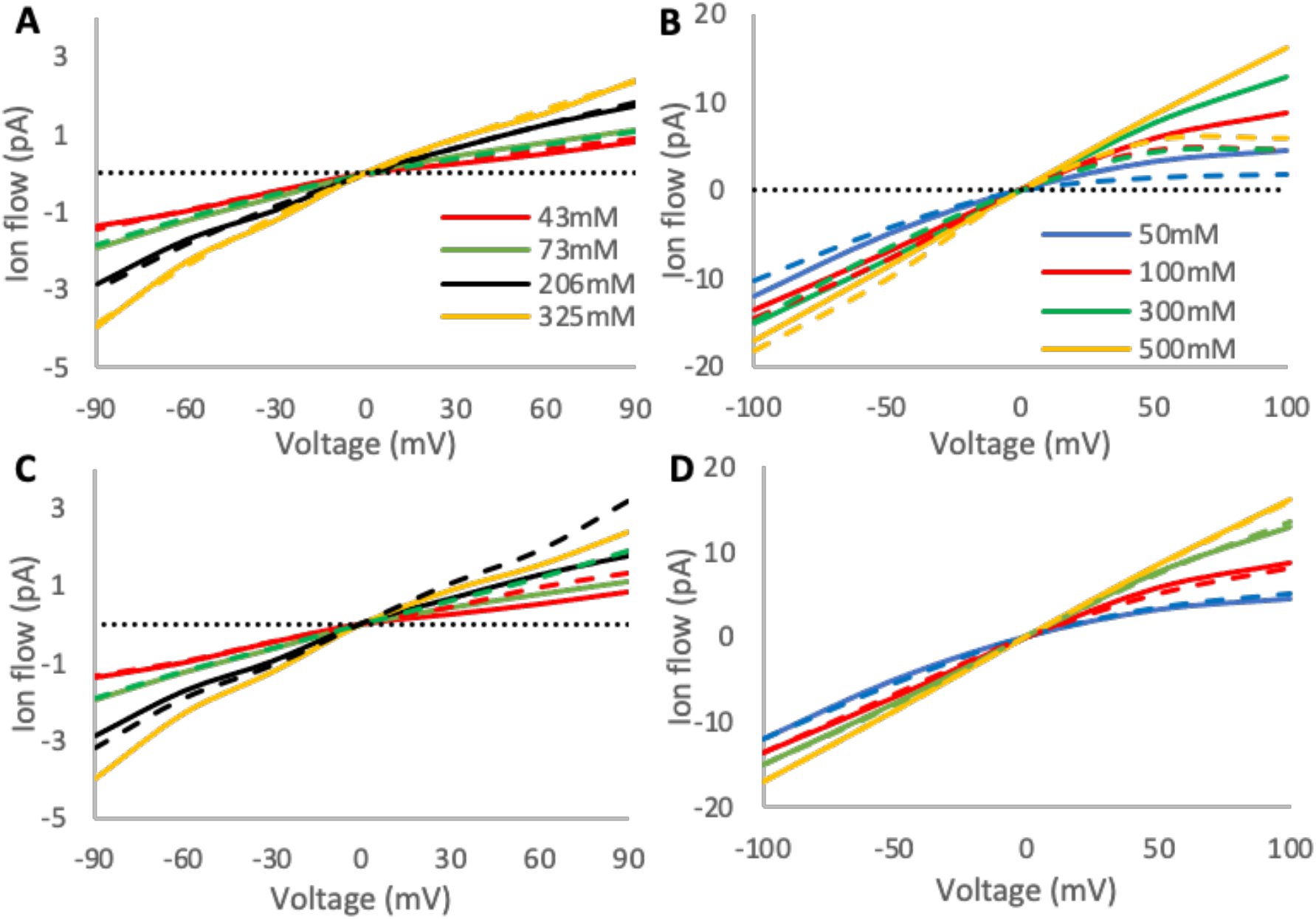
Single-channel I-V curves of WT Shaker^1^ and Shaker-P475D^2^. Solid lines represent experimental data; Dashed lines represent MsRKM results for 3-site models. (A) WT optimization. (B) P475D constrained optimization starting from WT seeds, only adjusting S1 binding, release, and S1→ S2 transfer rate coefficients. (C) Corresponding WT constrained optimization starting from mutant seeds and only adjusting S1 uptake, release, and S1→S2 transfer. D) P475D optimization.

A critical difference between the three-site MsRKM model and the previously proposed P475D Eyring model is the assumed impact of electrostatics across the entire pore region. The MsRKM model assumes that electrostatic changes at the channel mouth would not dramatically impact the free energy of transitions >17 Å away in the selectivity filter. Meanwhile, the Eyring model assumed the mutant would alter the stability of all the binding sites and barriers, including that of S4, which is ∼35 Å away. To test if the MsRKM could replicate the results of the Eyring model, we did an initial optimization of mutant experimental data using the same boundaries and acceptance criterion as the WT optimization. This optimization produced 95 (95%) mutant solutions that met the acceptance criterion. We repeated the secondary optimization using the 95 mutant solutions as seeds to test if the new mutant solutions could then back-map to reproduce the WT single-channel I-V curves. Again, only S1 uptake and release, and S1 → S2 transfer were allowed to change. None of the 95 mutants → WT optimizations produced a viable solution within the acceptance criterion (Figure 12C).

Thus, the 3-site MsRKM model could not find a solution that accurately describes both WT and mutant experimental data. This is distinct from the four-site model and suggests that limiting state space to only 3 binding sites is an oversimplification for the Shaker channel. Future work will explore the ramifications of additional sites and the optimal representation for Shaker. Regardless of the optimal representation, these results demonstrate that the number of binding sites directly impacts the model’s ability to replicate experimental data.

### 3.5 A Network Representation is Still Essential in Single-Pathway Dominant Mechanisms

Previous studies have demonstrated that non-dominant mechanistic pathways contribute to ion flux in ClC-ec1 and can eventually shift to be dominant under different electrochemical gradients. This suggests that kinetic selection in ClC-ec1 changes mechanistic pathways as a function of reaction conditions. However, understanding how the reaction network, including off-pathway and competing pathway flux, influences these dominant pathways remains relatively unknown. We initially assumed that off-pathway cycles would not impact the dominant pathway in Shaker since a single mechanistic cycle prevails under a wide range of experimental conditions (solution 2). However, when we pruned the network down to the dominant pathway alone (Figure 11A), we saw completely different model behavior (Figure 13A), indicating that the off-pathway flux remains essential in this solution.

**Figure 13.**
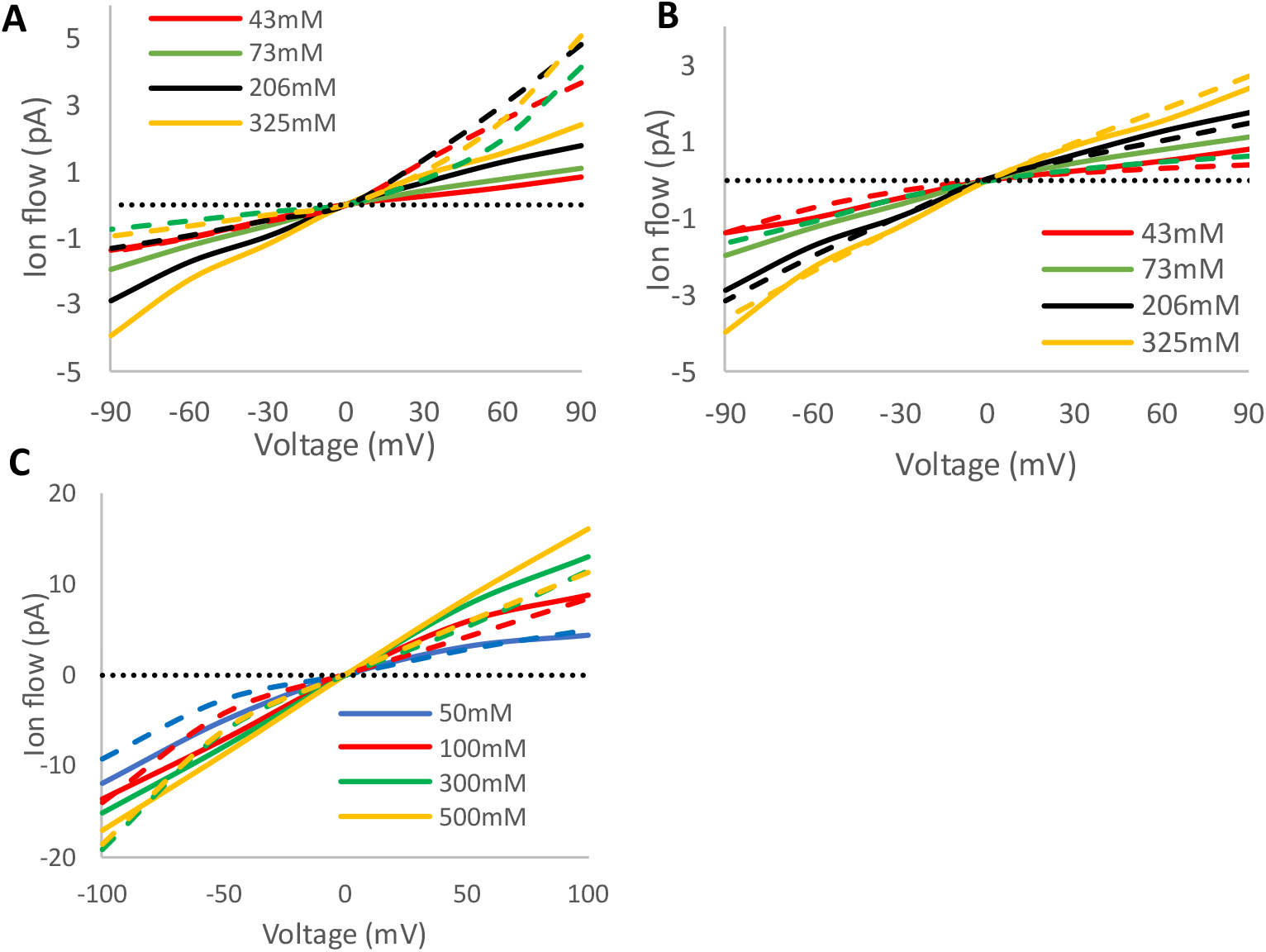
A) Single-channel I-V curves of WT^1^ (A, B) and P475D^2^ (C) Shaker channel. Solid lines represent experimental data; dashed lines represent MsRKM results for the dominant single pathway shown in 7A. A) WT network (solution 2). B) WT single pathway optimized. C) P475D mutant single pathway optimized

Since many published mechanisms of ion channels and secondary active transporters depict a single-cycle dominant mechanism, as seen for potassium channels^43, 45, 111^ and ClC-ec1,^27, 32, 112^ it is important to understand whether or not a network description of such mechanisms is important. Thus, to test if the contribution of off-pathway flux in solution 2 is simply an artifact of the MsRKM network solution refinement, we constructed a single-cycle kinetic model of the dominant pathway in solution 2 and optimized it to fit the WT experimental data using the same refinement methods used in the network implementation. However, the single-cycle kinetic model struggled to replicate the experimental data, deviating >0.3pA at every non-0 point along the I-V curve (Figure 13B). We then optimized the model to mutant data with a similar unconstrained optimization. Despite the mutant sharing the same dominant mechanistic pathway as the WT (Figure 11A, B), we observed significantly larger deviations (Figure 13C), potentially due to the larger flux observed in the mutant.

As noted in the three-site MsRKM, the number of intermediates strongly influences model behavior. Further, the single-cycle mechanism that we used was derived from the multi-pathway MsRKM solution 2, potentially biasing the system towards a multiple-cycle description. To remove these biases, we modeled the six-site mechanism described from MD simulations of the KcsA channel, a similar potassium channel^43^ (Figure 14). The fit of the six-site single-cycle model to the experimental data improves relative to the four-site single-cycle model, with the greatest single-point deviation just outside our acceptance criteria for the WT solution (Figure 14A). Notably, these deviations are still larger than the network optimized solution 2. The mutant six-site also performs better in an unconstrained mutant optimization (Figure 14B) compared to the four-site single-pathway model, but significant deviations remain compared to both experiment and the network optimized four-site model (Figure 11A).

**Figure 14.**
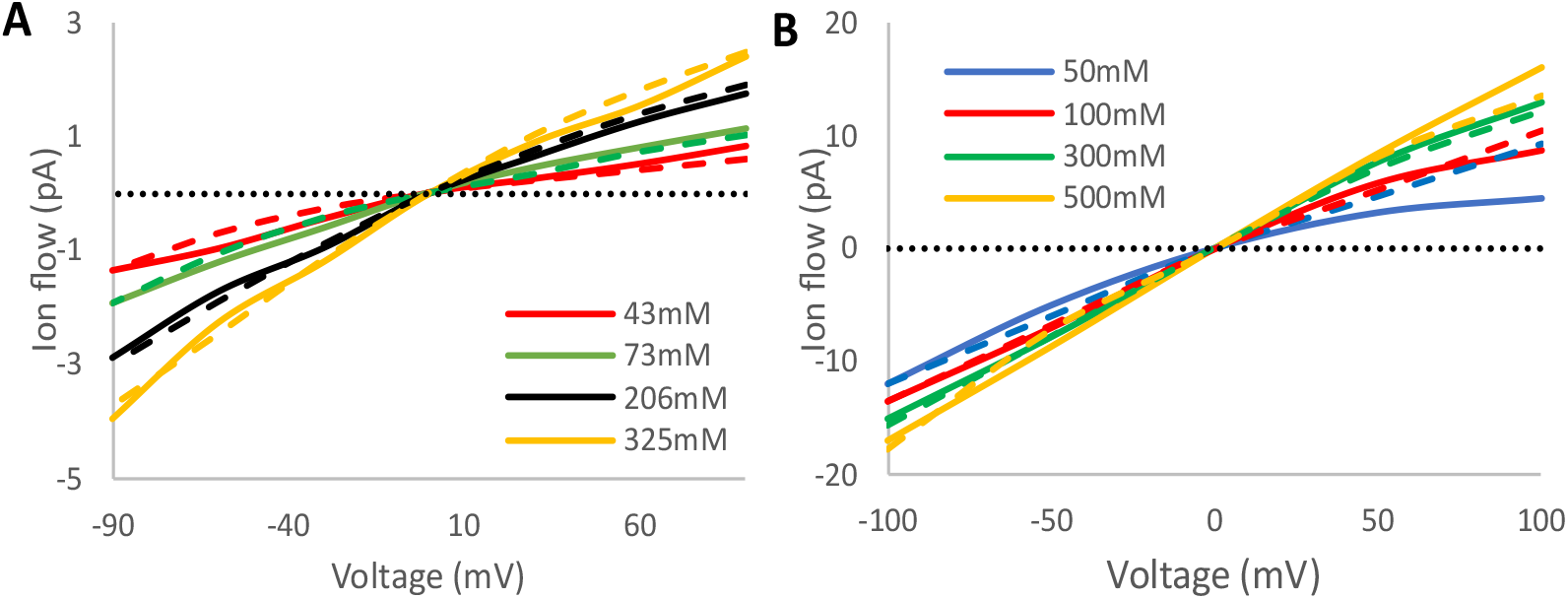
Single-channel I-V curves of WT (A) and P475D (B) Shaker channel. Solid lines represent experimental data; Dashed lines represent MsRKM results for optimized six-site MsRKM single-cycle models.

To further test the validity of a single-cycle representation, we examined the rectification properties of the P475D mutant, which change as a function of K^+^ concentration. Specifically, the degree of rectification decreases as concentration increases: the rectification ratio is ∼0.94 for +100mV/-100mV at 500 mM but drops to 0.36 for +100mV/-100mV at 50 mM. The WT data does not display this behavior, with rectification maintaining a ratio of roughly 0.60 for +90mV/-90mV from 43 mM to 325 mM. Surprisingly, none of the single-cycle models tested capture the changing rectification behavior of the P475D mutant. Instead, they maintain a rectification ratio close to that of the WT experimental data (∼0.64 for +/-100mV at all concentrations tested). The six-site, single-pathway mutant solution shows small changes in rectification as a function of concentration, which allows for a slightly better fit, but is still far off from the ∼0.58 rectification ratio change in the experimental data (Figure 14B). This problem was not observed for the multi-pathway model (Figure 12D), which implies that multiple mechanistic pathways may play an important role in concentration-dependent rectification. However, changing rectification as a function of concentration may be affected by site location and the dimensionless coupling factor. Future studies with a more detailed Shaker model will explore these relationships further. However, if the results found herein hold for more detailed models, a network representation that includes competing and off-pathway flux may be essential to describe concentration-dependent rectification properties.

## CONCLUSIONS

We have presented a new multiscale kinetic modeling framework that describes biomolecular processes involving multiple rare event transitions with reaction networks and the reactive flux through them under a range of equilibrium and non-equilibrium conditions. The framework was designed to combine the strengths of bottom-up rate quantification from multiscale simulations with top-down numerical solution refinement with experimental data to iteratively converge kinetic networks that not only describe known observables but are consistent with our best microscopic understanding. The obtained network descriptions can reveal when and how competing mechanistic pathways and off-pathway flux influence mechanistic outcomes. Motivated by the challenge of describing channels and transporters, the framework was designed to be responsive to electrochemical reaction conditions via voltage-dependent rate coefficients. This enables the use of and comparison to electrophysiology assays. Although MsRKM captures a system’s non-equilibrium response to a range of conditions, care should be taken to apply it to conditions for which the influence on underlying rate coefficients remains valid. In present form, this does not include significant conformational changes, such as those demonstrated in voltage-gated channels’ gating and inactivation.

The methods are demonstrated on the ClC-ec1 antiporter and the Shaker channel. For the more complex antiporter, ClC-ec1, determining a robust MsRKM solution requires integrating molecular-level data from simulations^6, 33, 34^ with ensemble data from experiments^26, 30, 31, 109^. Specifically, multiscale simulations provide estimated rate coefficients that help limit the vast solution space but require refinement based on experimental data. Increasing the variety in the experimental data further refines the solution space until robust and consistent solutions emerge. Importantly, the kinetic modeling predicted that certain rates require larger error bounds for low pH behavior than originally obtained in simulations. This prediction is consistent with recent cryoEM data demonstrating a pH-dependent conformational change.^29^ This is an example of how kinetic modeling can combine simulation and experimental data to iteratively converge on a consistent mechanistic description and to identify inconsistencies between the two.

The Shaker channel is modeled to test MsRKM in a purely top-down kinetic modeling approach based on experimental data. We demonstrate that robust solutions and physical insight can still be obtained in the absence of simulation-based rates. By iteratively increasing the integrated experimental data,^1, 2^ MsRKM converges to a solution that best predicts channel behavior, while revealing several interesting biophysical features. First, solutions that best fit WT Shaker and the P475D mutant had rate limiting steps before the selectivity filter, consistent with previous predictions^2^ for low-conductance K^+^ channels. Second, despite the inclusion of the entire reaction network, MsRKM converged on a single-pathway dominant mechanism (solution 2), verifying that the method can eliminate noise to identify single-pathway dominant processes. Third, the MsRKM process can reveal limitations in the state space definition. Reducing the four-site Shaker model to three binding sites failed to fit all known experimental data. Fourth, despite being dominated by a single pathway, the network representation, including competing and off-pathway flux, is still essential. The dominant pathway alone, with any rate definitions, fails to replicate experimental data. Even with the addition of binding sites and the use of a dominant pathway identified in simulations^43^, the six-site single-pathway model remains insufficient. In particular, single-pathway models fail to describe concentration-dependent rectification, suggesting that variable rectification ratios may require a network description. This is supported by the full four-site mutant solution, which revealed pathways sensitive to changing K^+^ and voltage gradients, resulting in variable rectification ratios.

While this paper focuses on channels and transporters, the MsRKM formalism is apt to treat any process involving multiple rare event transitions. It serves as a helpful bridge between experiment and simulation, enabling self-consistent data integration to identify and remedy inconsistencies. It additionally describes the non-equilibrium reaction flux that is often both biologically relevant and experimentally measured. This provides a consistent framework to describe and interpret a system’s equilibrium characterization and nonequilibrium behavior. Perhaps most importantly, it captures a complete picture of the reaction network and quantifies how competing mechanistic pathways collectively define the reactive flux and, thus, mechanism.

Future work will explore the incorporation rates responsive to pH-driven conformational changes. We will also test the hypotheses presented herein on more detailed Shaker models with more ion binding sites, the inclusion of competing water molecules, and system-specific voltage coupling factors.

## AUTHOR INFORMATION

### Notes

The authors declare no competing financial interest.

## ACKNOWLEDGMENT

The authors acknowledge Dimitry Vlachos and Benoit Roux for helpful discussions. This work was supported by NIH NIGMS (R35GM143117) and the computational resources provided by Expanse at the San Diego Supercomputing Center through the Advanced Cyberinfrastructure Coordination Ecosystem: Services and Support (ACCESS) program (allocation MCB200018) supported by NSF (grant nos. 2138259, 2138286, 2138307, 2137603, and 2138296), as well as the Center for High-Performance Computing (CHPC) at the University of Utah.

## For Table of Contents only

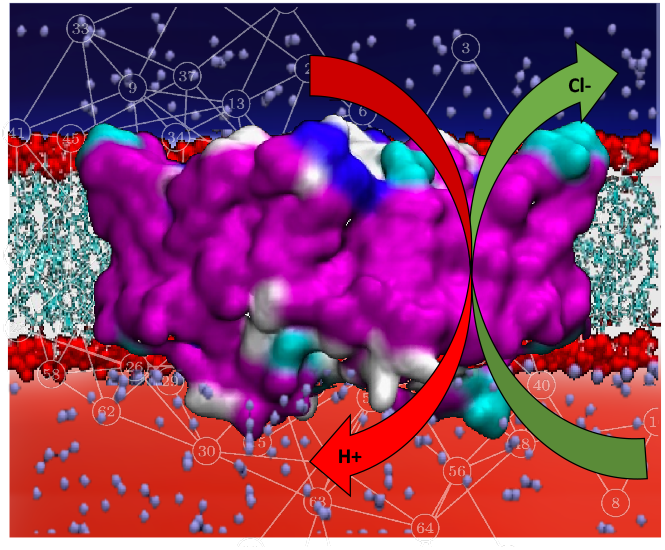

